# Kiaa1024L/Minar2 is essential for hearing by regulating cholesterol distribution in hair bundles

**DOI:** 10.1101/2022.06.23.497313

**Authors:** Ge Gao, Shuyu Guo, Quan Zhang, Hefei Zhang, Cuizhen Zhang, Gang Peng

## Abstract

Unbiased genetic screens implicated a number of uncharacterized genes in hearing loss, suggesting some biological processes required for auditory function remain unexplored. Loss of *Kiaa1024L*/*Minar2*, a previously understudied gene, caused deafness in mice, but how it functioned in hearing was unclear. Here we show that disruption of *kiaa1024L/minar2* causes hearing loss in the zebrafish. Defects in mechanotransduction, longer and thinner hair bundles, and enlarged apical lysosomes in hair cells are observed in *kiaa1024L/minar2* mutant. In cultured cells, Kiaa1024L/Minar2 is mainly localized to lysosomes and its overexpression recruits cholesterol and increases cholesterol labeling. Strikingly, an accessible pool of cholesterol is highly enriched in the hair bundle membrane, and loss of *kiaa1024L/minar2* reduces cholesterol localization to the hair bundles. Decreasing cholesterol levels aggravates, while increasing cholesterol levels rescues hair cell defects in *kiaa1024L/minar2* mutant. Therefore cholesterol plays an essential role in the hair bundles, and Kiaa1024L/Minar2 regulates cholesterol distribution and homeostasis to ensure normal hearing.

## Introduction

Hearing loss is one of the most common disabilities in human (World-Health-Organization, 2021). Studies of genes linked to non-syndromic deafness have identified over 120 genes essential for normal hearing (Shearer et al., 1993; Van Camp & Smith, 2021). This large collection of genes likely reflects the developmental and physiological complexity of the vertebrate auditory system. Large scale, unbiased genetic screens in the mouse (Bowl et al., 2017; Ingham et al., 2019) and the zebrafish (Whitfield et al., 1996) model systems have additionally identified multiple hearing loss genes. Interestingly, a number of these phenotypically identified hearing loss genes are previously uncharacterized and have no demonstrated functional roles, hinting that some biological processes required for normal auditory function have not been sufficiently explored (Bowl et al., 2017; Ingham et al., 2019).

The auditory hair cells are the sensory receptor cells that convert acoustic and mechanical stimuli into electrical signals initiating hearing (Fettiplace, 2017; Hudspeth, 1989; O. Maoileidigh & Ricci, 2019). Hair bundle, a specialized organelle situated at the apcial surface of each hair cell, responds to mechanical displacement in direction-dependent manner (Flock, 1964; Tilney et al., 1992). The hair bundle is essential for mechanoelectrial transduction (MET), and defects in the hair bundle can cause hearing loss (Belyantseva et al., 2009; Blanco-Sanchez et al., 2014; Kozlov et al., 2007; Noben-Trauth et al., 2003; Perrin et al., 2013). A number of proteins, such as adhension molecules, actin-bundling proteins, and disease-associated proteins are required for the morphogenesis and physiological regualtion of the hair bundle (Barr-Gillespie, 2015; Blanco-Sanchez et al., 2017; McGrath et al., 2017; Tilney, 1980). In addition, specific lipid molecules may play a role in the hair bundle. For instance, phosphatidylinositol-4,5-bisphosphate (PIP2) is localized to hair bundles, and it binds to the MET channel compment TMIE (Cunningham et al., 2020) and regulates the rates of fast and slow adaptation (Effertz et al., 2017; Hirono et al., 2004).

Cholesterol is an important component of eukaryotic cell membranes, and it controls membrane stiffness, tension, fluidity and other membrane properties (Maxfield & van Meer, 2010; Subczynski et al., 2017). Cholesterol also plays a regulatory function by interacting with membrane proteins (Harris, 2010). Previous studies showed that abnormally high or low cholesterol levels are detrimental to hearing (Corwin & Warchol, 1991; Crumling et al., 2012; Ding et al., 2020; Guo et al., 2005; King, Gordon-Salant, Pawlowski, et al., 2014; King, Gordon-Salant, Yanjanin, et al., 2014; Morizono, 1978; Sikora et al., 1986; Takahashi et al., 2016; Thoenes et al., 2015; Xing et al., 2015; Yao et al., 2019). Early studies also showed that cholesterol is not uniformly distributed in the hair cell membranes. The intensities of cholesterol labeling the hair cells were higher in the apical membranes when compared to those of the lateral membranes (Nguyen & Brownell, 1998; Takahashi et al., 2016), and freeze-fracture images of hair cells suggested that the stereocilia membrane was densely covered with cholesterols (Forge et al., 1988). Nevertheless, the distribution of cholesterol in hair cells in vivo, and cholesterol’s functional role in hair cell are not well characterized.

*Kiaa1024L*/*Minar2*, a previously understudied gene, was identified in a hearing loss screen in mouse knockout strains using the auditory brainstem response test. The homozygous *Kiaa1024L*/*Minar2* knockout mice aged 14weeks old had severely raised ABR thresholds at all frequencies tested (Bowl et al., 2017; Ingham et al., 2019), but how *Kiaa1024L*/*Minar2* functioned in hearing was unclear. Here we show *kiaa1024L/minar2* is expressed in the zebrafish mechanosensory hair cells, and disruption of *kiaa1024L/minar2* causes hearing loss in the zebrafish larvae. We next show that GFP-tagged Kiaa1024L/Minar2 protein is distributed in the stereocilia and the apical endo-membranes. Defects in mechanotransduction, longer and thinner hair bundles, and enlarged apical lysosomes are observed in hair cells in *kiaa1024L/minar2* mutant. In vitro studies in cultured cells show Kiaa1024L/Minar2 protein is mainly localized to lysosomes, and overexpression of Kiaa1024L/Minar2 recruits cholesterol and results in increased intracellular cholesterol levels. Strikingly, we show an accessible pool of cholesterol is highly enriched in the hair bundle membranes, and loss of *kiaa1024L/minar2* reduces cholesterol distribution in the hair bundles. Drug treatment decreasing cholesterol levels aggravates, whilst treatment increasing cholesterol levels rescues hair cell defects and hearing in the mutant *kiaa1024L/minar2* larvae. Together, our results indicate cholesterol plays an essential role in the hair-bundles, and Kiaa1024L/Minar2 regulates cholesterol distribution and homeostasis in auditory hair cells to ensure normal hearing.

## Results

### *kiaa1024L/minar2* is required for normal hearing in the zebrafish

*Kiaa1024L/minar2* gene orthologs are found in the vertebrate species only (***Figure 1-figure supplement 1A***). It belongs to a gene family that also includes *kiaa1024/minar1/ubtor* (Ho et al., 2018; Zhang et al., 2018). To conform to current human gene nomenclature, *kiaa1024L/minar2* gene orthologs are referred to as *minar2* hereafter.

We surveyed available sequencing data (Barta et al., 2018; Elkon et al., 2015; Erickson & Nicolson, 2015; Liu et al., 2018) and found the transcripts of *minar2* orthologs were highly enriched in the auditory hair cells of the mouse and the zebrafish (***Figure 1-figure supplement 1B***). In a human inner ear organoid model (Steinhart et al., 2022), *MINAR2* is specifically expressed in differentiated hair cells, similar to known differentiated hair cell markers (***Figure 1-figure supplement 1B***). Consistent with these sequencing based data, in situ hybridization results confirmed *minar2* was specifically expressed by the hair cells of the inner ears and the lateral line neuromasts in the developing zebrafish (***Figure 1A***).

**Figure 1.**
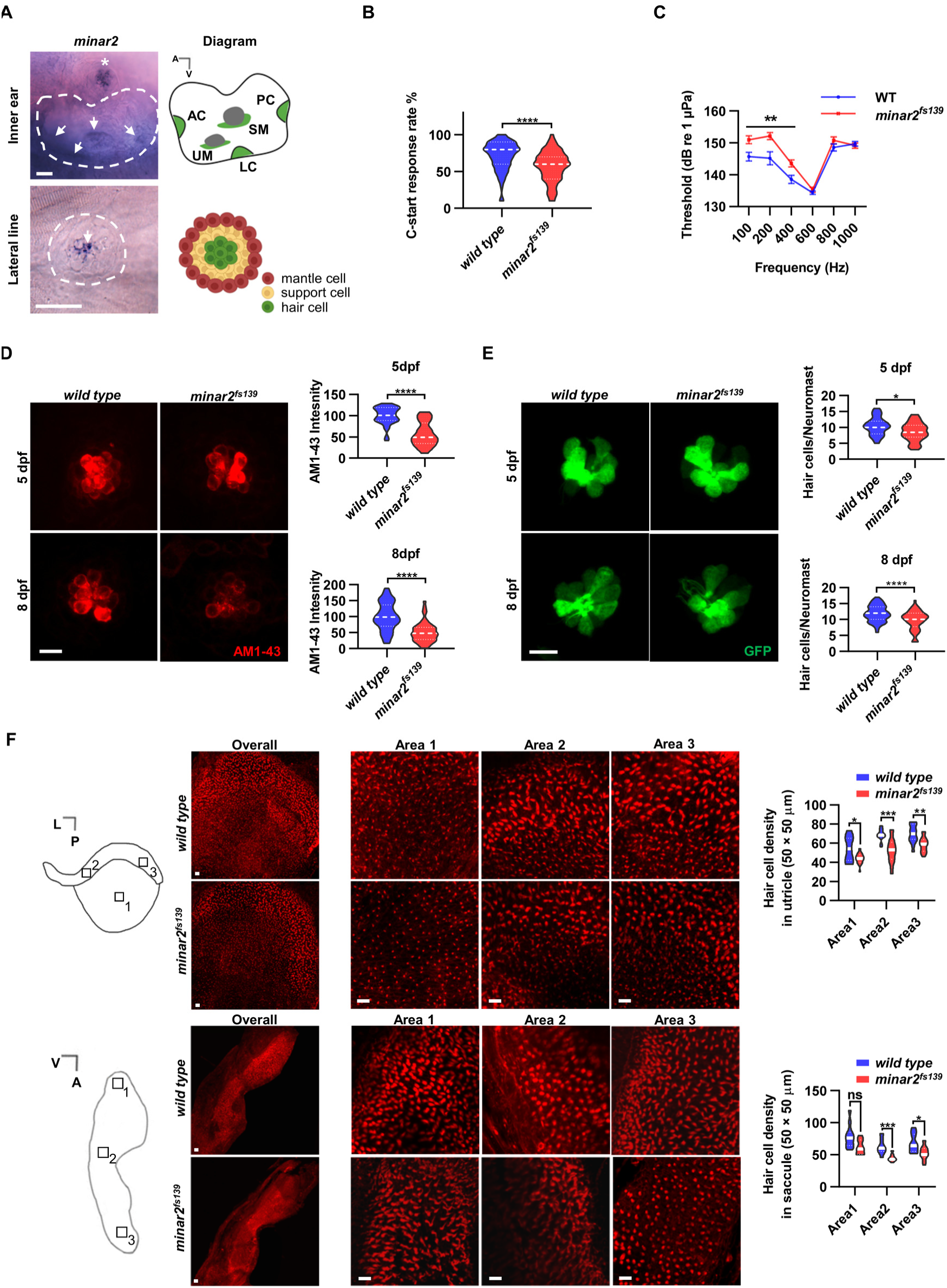
*minar2* is required for normal hearing in the zebrafish. **(A)** RNA in situ hybridization results showed *minar2* was specifically expressed by the hair cells of the inner ears and the lateral line neuromasts (5 dpf, days post fertilization). Arrows point to hair cells. Asterisk in the upper panel marks a head neuromast located next to the inner ear. AC: anterior crista; LC: lateral crista; PC: posterior crista; UM: utricular macula; SM: saccular macula. **(B)** C-start response rates for wild type and homozygous *minar2^fs139^* mutants at 8 dpf (n = 63 and 64, respectively. ****p < 0.0001, Mann-Whitney test). **(C)** Auditory evoked potentials (AEP) thresholds in wild type and the *minar2^fs139^* mutants (n = 11 and 13, respectively. For 100-, 200-, and 400-Hz, **p < 0.01). **(D)** Evaluation of mechanotransduction by AM 1–43 staining. The lateral line L3 neuromasts of 5 dpf and 8 dpf larvae were imaged and quantified (for 5 dpf, n = 35 and 36, t=7.465, df=64.84, ****p<0.0001; for 8 dpf, n = 49 and 51, t=6.444, df=86.90, ****p < 0.0001). **(E)** Quantification of hair cell numbers by counting the *myo6:Gal4FF;UAS-EGFP*-positive cells in lateral line L3 neuromast (for 5 dpf, n = 30 and 32, t=2.578, df=59.93, *p=0.0124; for 8 dpf, n = 58 and 58, t=4.148, df=114, ****p < 0.0001). **(F)** Quantification of hair cell numbers in the inner ears of zebafish adult. Hair bundles in dissected untricles (upper panels) and saccules (lower panels) were labeled with fluorescence-conjugated phalloidin. Diagrams of an untrile and saccule on the left. Numbered boxes (1-3) in the diagrams indicate the positions of imaged and counted areas (for utricles, n = 15 and 19; for saccules, n = 12 and 9. *p<0.05, **p<0.01, ***p<0.001). A: anterior; L: lateral; P: posterior; V: ventral. Scale bars represent 25 μm (A), and 10 μm (D, E and F). ***Figure 1-source data 1*** **Functional requirement and expression of *minar2* in hair cells** Figure 1B-1F; Figure 1-figure supplement 1B, 1D; Figure 1-figure supplement 2A-2C. Figure 1-source data 1.xlsx

To study *minar2* function in the zebrafish, we generated *minar2* mutant alleles by CRISPR/Cas9 mediated mutagenesis. The mutation in the *minar2^fs139^* allele was a 5-bp deletion in exon 1, and in the *minar2^fs140^* allele was a 5-bp insertion in exon 1 (***Figure 1-figure supplement 1C***). Both mutations were frameshift mutations and led to premature termination of protein translation. The translatable protein sequence in either mutant was very short and lacked the transmembrane helix located at the carboxyl terminus of the protein (***Figure 1-figure supplement 1C***). The mutant Minar2^fs139^ protein was translated to the 25th amino acid then adding 28 code-shifted residues before the reading frame stopped. The mutant Minar2^fs140^ protein was translated to the 26th amino acid then adding 55 code-shifted residues. Thus, both mutants were expected to be loss of function alleles. In addition, quantitative PCR analyses showed the expression levels of *minar2* in the two mutants were markedly down-regulated in the developing zebrafish, most likely because that the premature translational terminations activated the nonsense mediated mRNA decays (***Figure 1-figure supplement 1D***). We subsequently focused our investigation using the *minar2^fs139^* allele, and in some experiments corroborated our findings using the *minar2^fs140^* allele.

We first examined the short-latency C-start (SLC) response evoked by auditory stimuli (Burgess & Granato, 2007; Wolman et al., 2011) to assess the hearing abilities of zebrafish larvae. We found the SLC response rates to a 200-Hz stimulus were significantly reduced in the *minar2^fs139^* mutant at 8 dpf (***Figure 1B***, median response rates: 80 % in the wild type, and 60 % in the mutant. p < 0.0001, Mann-Whitney test).

To further analyze the hearing sensitivity of zebrafish larvae, we followed a procedure similar to the auditory brainstem response recording (Higgs et al., 2003; Higgs et al., 2002; Wang et al., 2015), and recorded auditory evoked potentials (AEP) in 7-8 dpf zebrafish larvae (***Figure 1C***). Two-way repeated-measures ANOVA revealed that the *minar2^fs139^* mutant had significantly elevated AEP thresholds compared with wild type control (For genotype factor, F(1, 22) = 7.457, p = 0.0122. For genotype x frequency, F(5, 110) = 6.072, p < 0.0001). The AEP thresholds were significantly higher at 100-400 Hz tone bursts in the *minar2^fs139^* mutant (for 100-Hz tone, 151.0 ± 1.2 dB for *minar2^fs139^*, 145.7 ± 1.4 dB for wild type control; for 200-Hz tone, 152.2 ± 1.1 dB for *minar2^fs139^*, 145.2 ± 2.0 dB for wild type control; for 400-Hz, 143.6 ± 1.1 dB for *minar2^fs139^*, 138.6 ± 1.3 dB for wild type control). These results indicate that *minar2* is required for normal hearing in the zebrafish larvae, and together with the loss of hearing phenotype in *Minar2* knockout mice (Bowl et al., 2017; Ingham et al., 2019), strengthen the conclusion that *minar2* orthologs play essential roles in the auditory functions of the vertebrates.

### Mechanotransduction are reduced in *minar2* mutant

Because *minar2* was specifically expressed by the mechanosensitive hair cells, we examined whether there were defects in the hair cells in the *minar2* mutants. We first determined levels of mechanotransduction in zebrafish larvae using AM1-43 labeling (***Figure 1D***). AM1-43, a fixable analog of FM1-43, is a vital fluorescence dye that labels hair cells by traversing open mechanosensitive channels (Meyers et al., 2003), thus providing an estimate of active mechanotransduction. We found the labeling intensities of AM1-43 were markedly reduced in the hair cells of the lateral line neuromasts of the *minar2^fs139^* mutant (56.8% and 51.1% of wild type control at 5 dpf and 8 dpf, respectively). We next counted the number of hair cells in the lateral line neuromasts by crossing the *minar2^fs139^* mutant with a *myo6:Gal4FF;UAS-EGFP* transgenic line, which specifically labeled hair cells (***Figure 1E***). The results showed the average numbers of hair cells were decreased in the *minar2^fs139^* mutant (82.3% and 82.1% of wild type control at 5 dpf and 8 dpf, respectively). Because the decreases of hair cell numbers (∼18% reduction) are smaller than those reductions of AM1-43 labeling intensities (∼50% reduction), these results suggest loss of *minar2* function mainly affect hair cell mechanotransduction in the zebrafish larvae.

We also examined the numbers of inner ear hair cells in the *minar2* mutant. In the zebrafish larvae, loss of *minar2* didn’t alter the number of hair cells in the inner ears (for 5 dpf wild type: 23.9 ± 0.6, *minar2^fs139^* mutant: 23.6 ± 0.4, *minar2^fs140^* mutant: 24.9 ± 0.6 hair cells, F(2, 82) = 1.399, p=0.253; for 8 dpf wild type: 32.5 ± 0.7, *minar2^fs139^* mutant: 32.9 ± 0.5, *minar2^fs140^* mutant: 33.0 ± 0.7 hair cells, F(2, 79) = 0.2378, p=0.789, both of the lateral crista of inner ear, ***Figure 1-figure supplement 2A***). The homozygous *minar2* mutants were viable, and the body length and body weight of adults were no different from the wild type controls (***Figure 1-figure supplement 2B***). Nevertheless, hematoxylin and eosin staining of head sections suggested the inner ear regions were smaller and the numbers of hair cells were decreased in utricle, semicircular canal crista, and saccule of the *minar2^fs139^* mutant (***Figure 1-figure supplement 2C***). To quantify the numbers of hair cells, we excised utricles and saccules from inner ears and labeled the hair cells with fluorescence-conjugated phalloidin. The average numbers of hair cells were broadly decreased across different regions of utricles and saccules in the *minar2^fs139^* mutant, ranging from 71.2% to 84.3% of wild type controls (***Figure 1F***, for the utricles, the genotype factor, F(1, 32) = 37.77, p<0.0001; for the saccules, the genotype factor, F(1, 19) = 19.94, p<0.001).

We conclude that *minar2* is required for mechanotransduction in the zebrafish larvae, and loss of *minar2* reduces the number of inner ear hair cells in the zebrafish adults.

### Minar2 protein localizes to the stereocilia and the apical region of the hair cells

To study how Minar2 functioned in the hair cells, we first examined the subcellular localization of Minar2 protein in the hair cells. As the Minar2 antibodies were not available, we generated a GFP-Minar2 fusion construct driven by a hair cell-specific promoter, *myosin 6b* (Kindt et al., 2012; Maeda et al., 2017). We injected this construct into fertilized oocytes and expressed the GFP-Minar2 fusion in the hair cells. We observed that GFP-Minar2 localized to the apical region of hair cell, apparently around and just below the cuticular plate. The GFP-Minar2 fluorescence also co-localized with the phalloidin labeled stereocilia (***Figure 2A***). These spatial distribution patterns of GFP-Minar2 were similar regardless of apparent expression levels. We subsequently generated a stable transgenic line using the *myo6b*:GFP-Minar2 construct and we found GFP-Minar2 also localized to the stereocilia and the apical region of the hair cells in the transgenic animals (***Figure 2B***). Prominent vesicle-like structures were seen below the cuticular plate, while a ring-like structure was often observed at the base of the hair bundle. We also co-labeled the kinocilia with anti-acetylated tubulin antibodies in the *myo6b*:GFP-Minar2 transgenic zebrafish and found there was no observable GFP-Minar2 signal in the kinocilia (***Figure 2-figure supplement 1A***).

**Figure 2.**
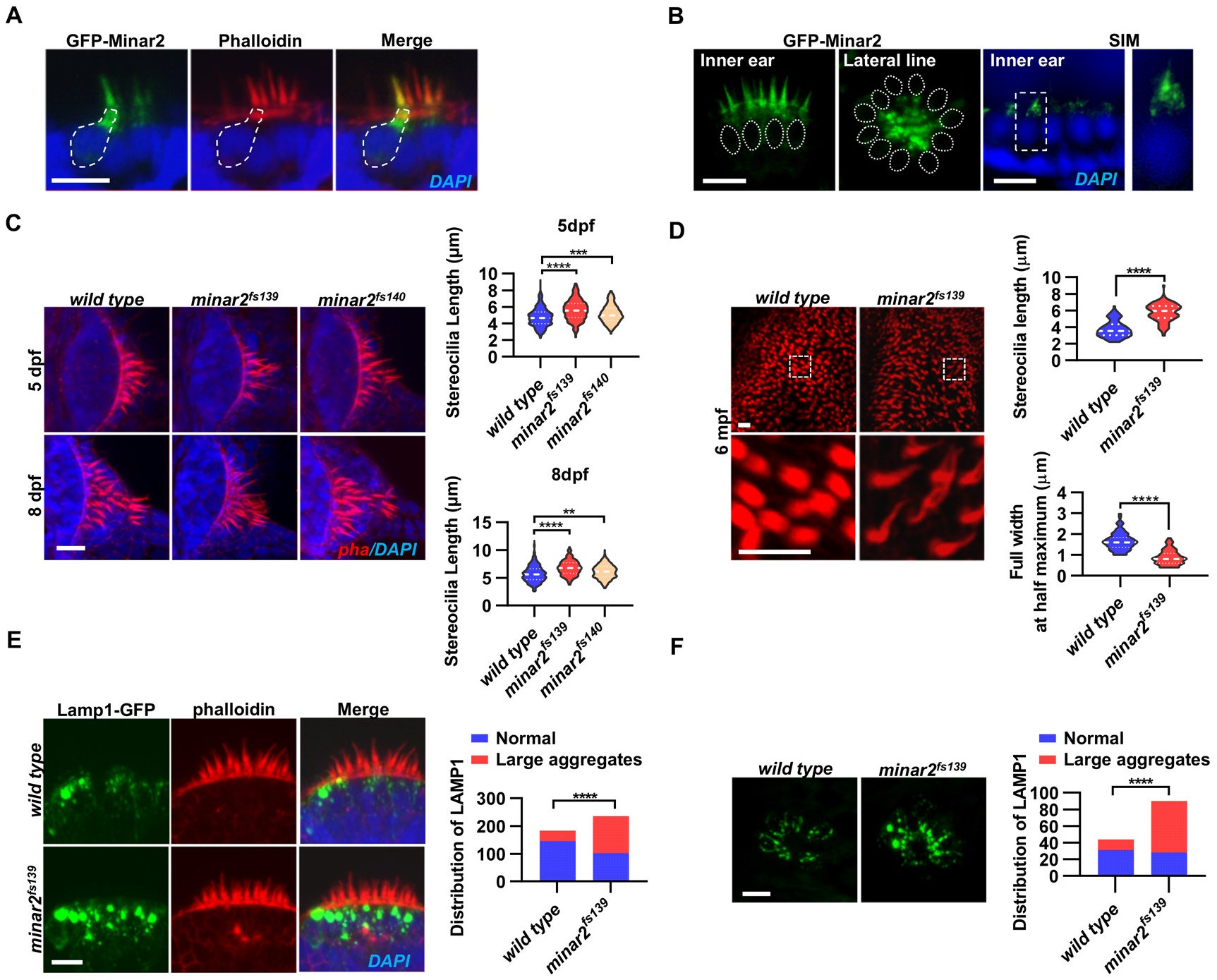
Localization and function of Minar2 in the stereocilia and the apical region of the hair cells. **(A)** Representative images of transiently expressed GFP-Minar2 in hair cells. Dashed line marks the border of a hair cell expressing GFP-Minar2. Stereocilia were labled with phalloidin. Nuclei were counter stained by DAPI. **(B)** Distribution of GFP-Minar2 in hair cells in the stable *myo6*:GFP-Minar2 transgenic line. Representative images of hair cells of lateral crista of inner ear (Inner ear) and lateral line neuromast (Lateral line). Dashed lines marks the nuclei of hair cells. Hair cells were also imaged with structured illumination microscopy (SIM), a super-resolution method. Right panel shows an enlarged view of the boxed area. **(C)** Quantification of hair bundlel lengths of the inner ear hair cells in zebrafish larvae. Hair bundles were labeled with phalloidin and the lateral crista regions of inner ears were imaged. Hair bundle lengths were measured from 34, 29, and 22 images of wild type, *minar2^fs139^*, and *minar2^fs140^* larvae at 5 dpf, or 36, 23, 23 images of respective larvae at 8 dpf. For 5 dpf, n = 340, 290, and 220, F(2, 847) = 42.58, p<0.001; For 8 dpf, n = 360, 230, and 230, F(2, 817) = 42.95, p<0.001. Multiple comparison significance values are indicated on the graph. **(D)** Quantification of hair bundlel lengths and width of inner ear hair cells in zebrafish adult (6 mpf). Bottom panels show enlarged views of the boxed area. Hair bundles in the saccules were measured from 8 images for the wild type, and 8 images for the *minar2^fs139^* mutant. n = 80 and 80. ****p<0.0001. **(E-F)** Morphology and distribution of Lamp1-labeled lysosomes in the hair cells of innear ear (E) and lateral line neuromast (F) in zebrafish larvae (5 dpf). For inner ear, 36 and 44 images of lateral crista regions in the wild type and *minar2^fs139^* mutant were counted, respectively (n = 184 and 236, ****p<0.0001, Fisher’s exact test). For lateral line, 10 and 15 images of lateral line L3 neuromasts were counted (n = 44 and 90, ****p<0.0001, Fisher’s exact test). Scale bars represent 10 μm. ***Figure 2-source data 1*** **Localization and function of Minar2 in the apical regions of hair cells** Figure 2C-2F. Figure 2-source data 1.xlsx

To better examine the subcellular localization, we imaged hair cells with structured illumination microscopy (SIM), a super-resolution microscopy method. The SIM results showed that GFP-Minar2 was broadly distributed in the stereocilia. Below the stereocilia and in the apical region of the hair cell, GFP-Minar2 signal was composed of multiple small sized vesicles of various shapes (***Figure 2B***).

### Hair bundles are longer and thinner in *minar2* mutant

Because hair bundles are essential for mechanotransduction, and Minar2 protein localizes to the hair bundles and the apical regions where the hair bundles reside, we next examined the hair bundles in the *minar2* mutant. We labeled the hair bundles with phalloidin and observed that the hair bundles of inner ear hair cells were longer in the mutant *minar2^fs139^* and *minar2^fs140^* larvae (***Figure 2C***, for 5 dpf, wt: 4.77 ± 0.06 μm, *minar2^fs139^*: 5.62 ± 0.07 μm, *minar2^fs140^*: 5.15 ± 0.07 μm, F(2, 847) = 42.58, p<0.001; for 8 dpf, wt: 5.72 ± 0.08 μm, *minar2^fs139^*: 6.80 ± 0.09 μm, *minar2^fs140^*: 6.10 ± 0.08 μm, F(2, 817) = 42.95, p<0.001), and the hair bundles also appeared thinner. The thinning and lengthening of the mutant hair bundles were more pronounced in adult *minar2^fs139^* mutant (***Figure 2D***). In the 6-month-old adults, the average length of mutant hair bundles was more than 50% longer that of wild type controls (wt: 3.77 ± 0.11 μm, *minar2^fs139^*: 5.77 ± 0.12 μm, t = 12.05, df= 158, p<0.0001), and the width was only half of the controls (wt: 1.66 ± 0.05 μm, *minar2^fs139^*: 0.88 ± 0.04 μm, t = 12.65, df = 158, p<0.0001).

### Enlarged lysosome aggregates locate at the apical region of the hair cells in *minar2* mutant

Previous electric microscopy and immunofluorescence microscopy studies showed that the apical region of the hair cell is teemed with endocytotic vesicles, many of which are lysosomes (Revelo et al., 2014; Spicer et al., 1999; Wiwatpanit et al., 2018). We generated a GFP fusion construct for Lamp1, a lysosome marker, and expressed the Lamp1-GFP fusion in the hair cells using the *myo6b* promoter. Similar to the findings in the mouse model, we observed that Lamp1-GFP positive lysosomes were abundantly distributed in the apical region of the inner ear hair cells. There were also a few lysosomes distributed along the basolateral membrane of the hair cells. In the wild type controls, most of these Lamp1-GFP labeled lysosomes were small sphere- or rod-shaped vesicles. Occasionally, rare and large-sized Lamp1-GFP labeled structures were observed (in 39 out of 184 hair cells) and these structures likely were abnormally enlarged lysosomes or lysosome aggregates (diameter > 2 μm). In contrast, in the inner ear hair cells of the *minar2^fs139^* mutant, the large round-shaped lysosomal structures (diameter > 2 μm) were frequently observed (observed in 134 out 236 hair cells), and these abnormally enlarged lysosome aggregates were always located at the apical region of the hair cells (***Figure 2E***, p<0.0001, Fisher’s exact test). Similar results showing abnormally enlarged lysosome aggregates were also observed in the hair cells of the lateral line neuromasts (***Figure 2F***, p<0.0001, Fisher’s exact test). These data suggested that Minar2 may play a role in regulating the apical lysosomes in the hair cells. Consistent with this view, GFP-Minar2 signals partially overlapped with the Lamp1-mCherry signals at the apical region of the inner ear hair cells (***Figure 2-figure supplement 1B***).

We next examined the subcellular localization of Minar2 in cultured cells in vitro. Because *MINAR2* gene is expressed at low levels in cultured human cell lines (<1.4 nTPM in over 60 human tissue cell lines, Human Protein Atlas, proteinatlas.org), we expressed a GFP-Minar2 fusion construct in cultured HEK293 and Cos-7 cells, and used KDEL, GCC1/GM130, and Lyso-Tracker to label endoplasmic reticulum, Golgi complex, and lysosome respectively. We found GFP-Minar2 most strongly co-localized with lysosomes (***Figure 2-figure supplement 1C-1E***). Furthermore, when the morphology and distribution of the lysosomes were altered after treatment with U18666A, GFP-Minar2 signals followed the changes of lysosomes, increased in the particle sizes and accumulated towards the nuclear region (***Figure 2-figure supplement 1E***). We further stained the lysosome lumen with filipin staining of cholesterol, which was trapped inside lysosome after treatment with U18666A; and we observed that GFP-Minar2 signals appeared as circles that circumscribed the filipin signals, indicating GFP-Minar2 was localized on the lysosomal membranes (***Figure 2-figure supplement 1F***).

### Minar2 increases cholesterol labeling and colicalizes with cholesterol in cultured cells

Because exhaustive protein sequence homology searches failed to identify any funcitional domains in Minar2, we attempted sequence pattern searches (Liu et al., 2006) to provide clues into Minar2’s function. We first extracted highly conserved sequence patterns from the multiple sequence alignment result of Minar2 protein orthologs (***Figure 1-figure supplement 1A***), then performed protein pattern search against the Swiss-Prot database. One of the conserved sequence patterns (π-S/T-Ω-S/T-Ψ-ζ-ζ-Ω) had 125 hits in the Metazoa [taxid:33208] proteins, and 37 out of the 125 hit sequences matched to Caveolin-1 protein from various species (***Figure 3A***). A close inspection revealed that the matched sequence pattern resided in the caveolin scaffolding domain (CSD), which is known to interact with other proteins (Murata et al., 1995; Razani et al., 2002) and lipid membranes (Schlegel et al., 1999). The CSD of caveolin also directly binds to cholesterol in the membranes (Ikonen et al., 2004; Liu et al., 2016; Murata et al., 1995) and this binding may contribute to enrichment of cholesterol in the caveolae (Everson & Smart, 2005; Frank et al., 2006).

**Figure 3.**
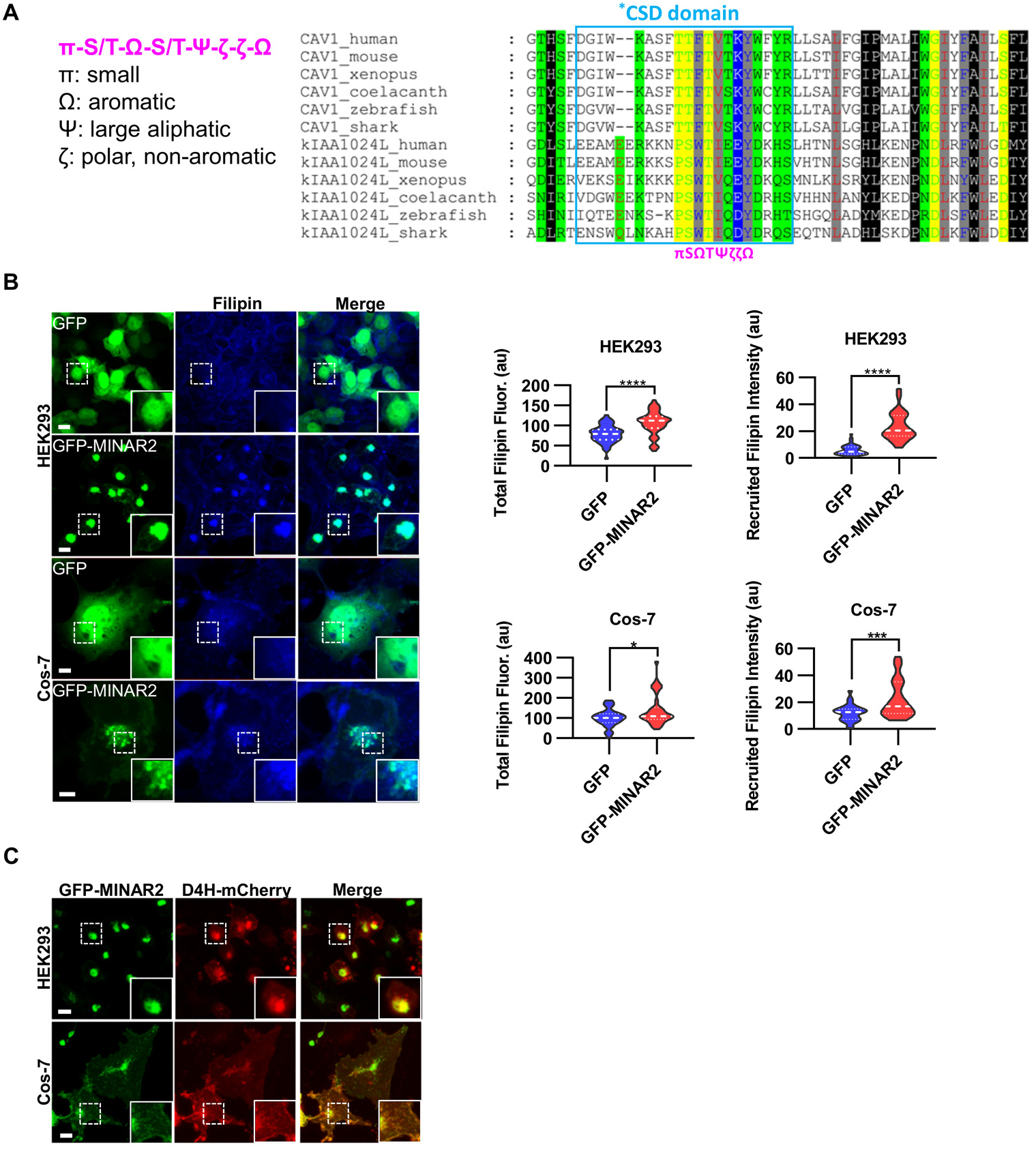
Minar2 increases cholesterol labeling and colicalizes with accessible cholesterol in cultured cells. **(A)** Protein sequence pattern search for Minar2 identifies caveolin. The conserved Minar2 sequence pattern is written in normalized symbols (Aasland et al., 2002; Livingstone & Barton, 1993). Sequence alignment is highlighed by physico-chemical properties of the amino acids. *CSD: caveolin scaffolding domain. **(B)** Effects of Minar2 on levels and distributions of filipin labeling in cultured cells. Total filipin fluorescence indicates sum of all pixel values of filipin signals. Recruited filipin represents average of pixel values of filipin signals located within GFP-positive area. For HEK293 cells, n = 51 and 58; for Cos-7 cells, n = 28 and 35. au: arbitory unit. **(C)** Distriubtion of GFP-MINAR2 and D4H-mCherry in cultured cells. Figure inserts show enlarged views of the boxed area. Scale bars represent 10 μm. ***Figure 3-source data 1*** **Effects of Minar2 on levels and distributions of cholesterol in cultured cells** Figure 3B. Figure 3-source data1.xlsx

We next investigated if Minar2 may function through cholesterol. We expressed the GFP-Minar2 fusion construct in HEK293 and Cos-7 cells, then stained these cells with filipin, a fluorescent polyene antibiotic that recognizing unesterified cholesterol in biological membranes (Maxfield & Wustner, 2012; Severs, 1997). In the control cells expressing GFP, the filipin signals were diffusely distributed across the cells and there was no specific overlap between the GFP and the filipin signals. In contrast, in cells expressing GFP-Minar2 fusion protein, the filipin signals were accumulated to the perinuclear regions and there were strong overlaps between the Minar2 and the filipin signals in the perinuclear regions (***Figure 3B***). Quantification of the filipin signals revealed that there were significant increases of intracellular unesterified cholesterol in cells expressing GFP-Minar2 (total filipin fluorescence, for HEK293 cells, 79.0 ± 3.0 au for GFP-only control, 106 ± 3.8 au for GFP-Minar2 group, t=5.654, df=104.6, p<0.0001; for Cos-7 cells, 100.5 ± 8.2 au for GFP-only control, 140.3 ± 12.9 au for GFP-Minar2 group, t=2.608, df=55.73, p=0.012). The quantification results also confirmed that GFP-Minar2 recruited the filipin signals to where GFP signals were located (recruited filipin intensity, for HEK293 cells, 5.85 ± 0.51 au for GFP-only control, 23.56 ± 1.36 au for GFP-Minar2 group, t=12.20, df=72.42, p<0.0001; for Cos-7 cells, 12.08 ± 1.1 au for GFP-only control, 22.14 ± 2.31 au for GFP-Minar2 group, t=3.903, df=48.96, p<0.001).

Previous studies suggested that cholesterol in the biological membranes is organized in three pools, a fixed pool for membrane integrity, a sequestered pool with low chemical activity, and an active/accessible pool with high activity (Das et al., 2014; Lange & Steck, 2016; Lange et al., 2004). Protein probes derived from bacterial toxin Perfringolysin O (PFO) and the minimal cholesterol-binding domain 4 (D4 or D4H) of PFO have been widely used to measure the accessible pool of cholesterol in biological membranes (Das et al., 2014; Lim et al., 2019; Maekawa & Fairn, 2015; Naito et al., 2019; Schoop et al., 2021). We expressed a D4H-mCherry probe to examine effects of Minar2 on the accessible cholesterol in cultured cells. Since the D4H-mCherry protein probe was expressed in the cytoplasm, and unlike filipin the protein probe cannot cross membranes, D4H-mCherry only recognizes accessible cholesterol on the cytosolic leaflets of plasma and organelle membranes. Similar to what observed in cells stained by filipin, GFP-Minar2 co-localized with D4H-mCherry signals, and recruited the D4H-mCherry signals to the perinuclear regions (***Figure 3C***).

We conclude that Minar2 co-localizes with assessable pool of cholesterol, and Minar2 may increases intracellular cholesterol levels when overexpressed in vitro.

### The hair bundle membrane is enriched for accessible cholesterol

Previous studies showed that cholesterol is not uniformly distributed in the hair cell membranes. In isolated outer hair cells and in outer hair cells in fixed cochlear tissues, cholesterol levels are higher in the in the apical membranes than those in the lateral wall membranes (Nguyen & Brownell, 1998; Takahashi et al., 2016).

To examine cholesterol distribution in the hair cells in zebrafish, we first performed filipin staining on fixed zebrafish tissues. The apical regions of inner ear hair cells showed distinct filipin labeling (***Figure 4-figure supplement 1A***). The signal-noise ratios of filipin labeling were not high, similar to results of filipin labeling on fixed cochlear tissues from mice (Rajagopalan et al., 2007). This is likely because that filipin stains unesterified cholesterol in all membranous structures, and that filipin is not very stable and photo-bleaches rapidly, making it less suitable on fixed tissue samples.

We next investigated the intracellular distribution of cholesterol in vivo by taking advantage of the D4H-mCherry protein probe. Strikingly, when expressed in the inner ear hair cells by microinjection, D4H-mCherry specifically labeled the stereocilia regardless of its expression levels (***Figure 4A***). Low-intensity labeling of kinocilia was also observed in cells with high level of D4H-mCherry expression, while the basolateral membranes of hair cells were not or only very faintly labeled (***Figure 4A***). To exclude possible compounding factors of expression levels of probes or special membrane structures of stereocilia, we co-expressed PM-GFP, a membrane probe containing the N-terminal 10 amino acid sequence of Lyn sufficient for palmitoylation and myristoylation modification and plasma membrane targeting (Pyenta et al., 2001). As expected, PM-GFP uniformly labeled the basolateral membranes, together with the stereocilia and the kinocilia membranes (***Figure 4A***). In the hair cells of the later line neuromasts, the stereocilia were similarly strongly labeled by D4H-mCherry. There were also observable but weaker D4H-mCherry signals on the basolateral membranes of the later line hair cells (***Figure 4A***). These data suggested that there are high levels of accessible cholesterol located to the stereocilia membranes, while the accessible cholesterol levels are markedly lower in the basolateral membranes in the hair cells.

**Figure 4.**
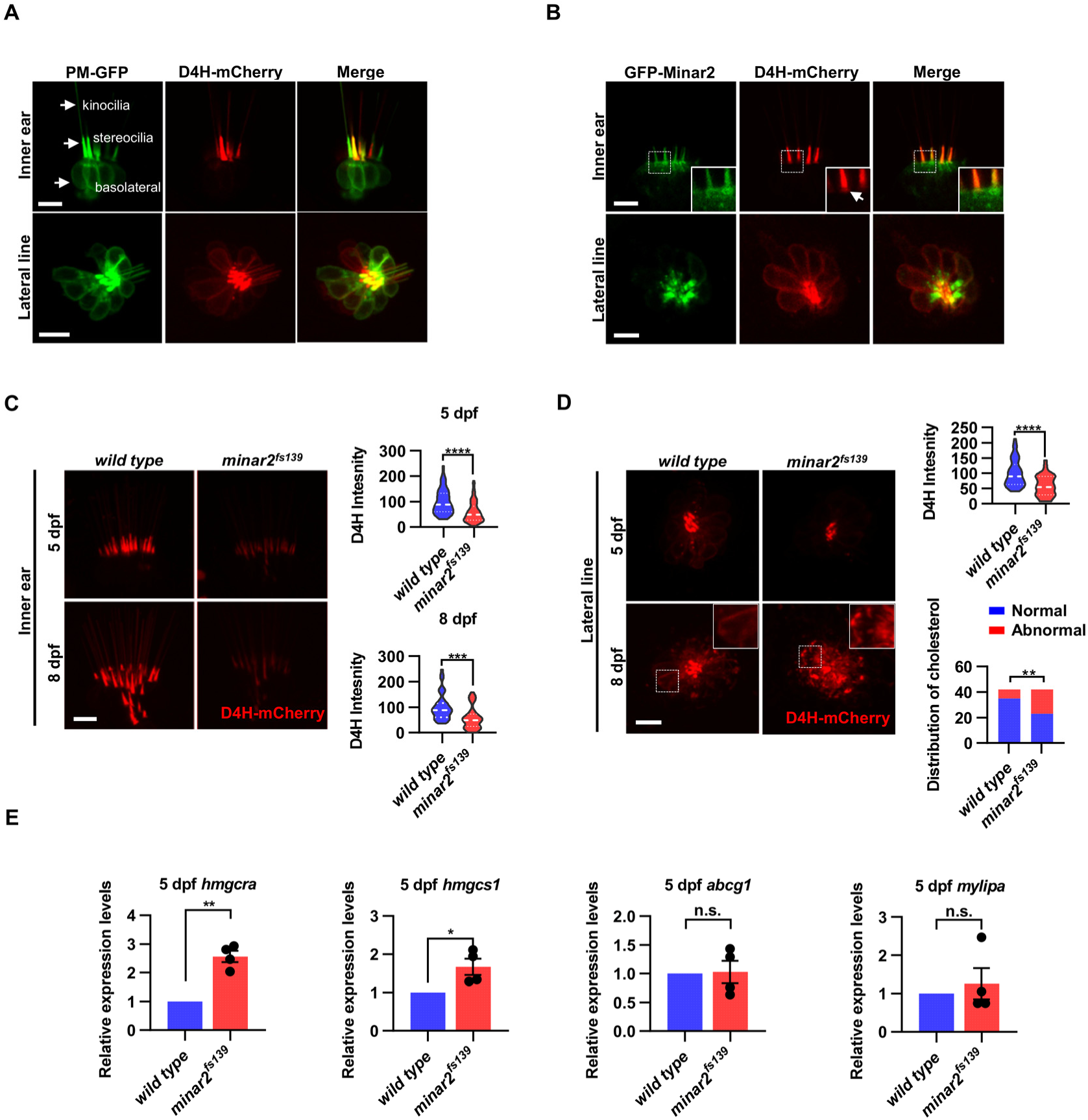
Cholesterol labeling in the stereocilia is reduced in *minar2* mutant. **(A)** Representative images of PM-GFP and D4H-mCherry expressed in hair cells. The general membrane probe PM-GFP labels kinocilia, stereocilia and basolateral membranes (arrows). The accesible cholesterol probe D4H-mCherry mostly labels the stereocilia in the inner ear. The lasteral crista of inner ear and the L3 lateral line neuromast were imaged. **(B)** Distribution of GFP-Minar2 and D4H-mCherry in hair cells in stable transgenic zebrafish. GFP-Minar2 and D4H-mCherry entensively co-localize in the stereocilia, and in a few structures just below the stereocilia (arrow in figure insert). **(C)** Quntification of the intensity of cholesterol probe D4H-mCherry in the inner ear hair cells. The lateral crista regions of the inner ears were imaged and quantified. For the 5 dpf groups, n = 49 and 49 for the wild type and the *minar2^fs139^* mutant, repectively. t=4.446, df=93.30, ****p<0.0001; For the 8 dpf groups, n = 39 and 36, t=3.982, df=72.30, ***p<0.001. **(D)** Quntification of the intensity and appearance of D4H-mCherry in the lateral line hair cells. The lateral line L3 neuromasts were imaged and quantified. For the 5 dpf groups, the intensity of D4H-mCherry was quantified. n =40 and 33, t=4.438, df=70.81, ****p<0.0001. For the 8 dpf groups, the appearance of abnormally enlarged vesicles were quantified. Figure inserts show large vesicles in the basolateral regions in the *minar2^fs139^* mutant. n = 42 and 42, **p<0.01, Fisher’s exact test. **(E)** Quantification of expression levels of genes involved in cholesterol metabolism. Srebp2 target gene (*hmgcra* and *hmgcs1) and* LXR target gene *(abcg1* and *mylipa)* were examined by qRT-PCR. Expression levels relative to GAPDH levels were normalized to the wild type control group. For *hmgcra*, t=7.805, df=3, **p<0.01. For *hmgcs1*, t=3.217, df=3, *p<0.05. Scale bars represent 10 μm. ***Figure 4-source data 1*** **Effects of *minar2* loss-of-function on cholesterol in the aprical regions of hair cells** Figure 4C-E; Figure 4-figure supplement 1B. Figure 4-source data 1.zip

### Accessible cholesterol labeling in the stereocilia is reduced in *kiaa1024L/minar2* mutant

Because both GFP-Minar2 and D4H-mCherry were distributed in the stereocilia (***Figure 2B*** and ***Figure 4A***), we asked if the two signals co-localized in the hair cells. We first generated a stable transgenic reporter line of the *myo6:D4H-mCherry* construct and crossed the D4H-mCherry report line to the stable *myo6:GFP-Minar2* line. As expected, both GFP-Minar2 and D4H-mCherry signals clearly co-localized in the stereocilia (***Figure 4B***). Just below the stereocilia and in the apical region of hair cells, there were abundant GFP-Minar2 signals, and close inspections revealed a few D4H-mCherry labeled structures that co-localized with GFP-Minar2 (***Figure 4B*** insert). D4H-mCherry additionally labeled kinocilia, in which GFP-Minar2 signals were absent. Thus, similar to co-localization results in cultured cell in vitro (***Figure 3C***), Minar2 co-localized with accessible cholesterol in the stereocilia in vivo.

We next examined the status of accessible cholesterol in the *minar2^fs139^* mutant by crossing the D4H-mCherry report line to the mutant *minar2^fs139^* line. The D4H-mCherry report line had a single transgenic insertion (***Figure 4-figure supplement 1B***), and we set the crossing scheme so that the wild control and the homozygous mutant larvae both had one copy of the D4H-mCherry transgene. We then analyzed the D4H-mCherry labeling in inner ear and neuromast whole mounts at 5 and 8 dpf. In the hair cells of inner ears in the mutant *minar2^fs139^* larvae, D4H-mCherry still labeled stereocilia (***Figure 4C***), but the intensities of the signals were dramatically reduced. To quantify and compare the fluorescence signals, we used custom MATLAB scripts to count pixel intensity values using optical sections through the hair cells. The results showed the average D4H-mCherry signals of inner ear stereocilia in the *minar2^fs139^* larvae were about half of those wild controls (59.16 ± 5.92% and 56.62 ± 7.15% for 5 and 8 dpf, respectively). The decreases of D4H-mCherry signals were not due to reduction of hair cell numbers, since there were no significant differences in the numbers of inner ear hairs at these developmental stages (***Figure 1-figure supplement 2A***). The decreases were not due to changes of transgene expression either, since the Western blotting results showed that the expression levels of D4H-mCherry were similar between the wild type controls and the mutant *minar2^fs139^* larvae (***Figure 4-figure supplement 1B***).

In the hair cells of lateral line neuromasts, there was a similar reduction of D4H-mCherry signal at 5 dpf (59.30 ± 5.98% of controls), while the signal intensity was no different from the wild type controls at 8 dpf (121.39% of controls, n = 41 and 44 for the wild type and the *minar2^fs139^* mutant, repectively. t=0.8889, df=83, p=0.3766). The discordancy between 5 and 8 dpf was also evident in the appearances of the D4H-mCherry signals. While D4H-mCherry primarily labeled the stereocilia membranes in both wild type controls and the *minar2^fs139^* mutant at 5 dpf, D4H-mCherry labeled many large, round- and rod-shaped vesicles in the cell body regions of the neuromast hair cells in the *minar2^fs139^* mutant at 8 dpf (***Figure 4D***, p<0.01, Fisher’s exact test). These D4H-mCherry labeled large vesicles in the hair cells of *minar2^fs139^* mutant likely corresponded to enlarged lysosomes, as these abnormal vesicles were co-labeled by the lysosome probe Lamp1-GFP, but not by the plasma membrane probe PM-GFP (***Figure 4-figure supplement 1C***).

Because the accessible cholesterol pool has high activity biochemically (Lange & Steck, 2016; Lange et al., 2004; Lim et al., 2019), we wondered whether the reduction of accessible cholesterol in the *minar2^fs139^* mutant may have consequences for the cholesterol metabolism. We examined expressions of genes downstream of Srebp2 and LXR, two master regulators of sterol synthesis and turnover (Aqul et al., 2011; Liu et al., 2009). We found that there were significant increases of relative mRNA expression for Srebp2 target gene *hmgcr* (2.6 ± 0.2 fold) and *hmgcs1* (1.7 ± 0.2 fold) in the *minar2^fs139^* mutant, while expression levels of LXR target gene *abcg1* and *mylip* were not changed (***Figure 4E***). These results suggested that the sterol synthesis pathway was activated due to a reduction of accessible cholesterol pool in the *minar2^fs139^* mutant larvae.

We conclude that *minar2* is required for the normal cholesterol distribution and homeostasis in the hair cells.

### Lowering cholesterol levels aggravates hair cell defects in *minar2* mutant

To further examine the involvement of cholesterol in *minar2* loss of function, we manipulated intracellular cholesterol levels using pharmacological treatments. Since the hair cells of lateral line neuromasts are superficially located, making it straightforward to apply pharmacological agents, we focused our investigation on the neuromast hair cells. We first used 2-hydroxypropyl-β cyclodextrin (2HPβCD), an oligosaccharide and a cholesterol chelating reagent widely used to extract cholesterol from live cells (Kilsdonk et al., 1995; Zidovetzki & Levitan, 2007). As expected, treatment with 2HPβCD markedly decreased D4H-mCherry signals in the neuromast hair cells, indicating the cholesterol levels were reduced (16.5 ± 2.4% of those of control, ***Figure 5A***). Moreover, the AM1-43 labeling intensities were significantly reduced after the 2HPβCD treatment (33.5% of those of controls), indicating the mechanotransduction was compromised when intracellular cholesterol was extracted (***Figure 5B***). The AM1-43 labeling intensities in the 2HPβCD treatment group were clearly lower than those in the *minar2^fs139^* mutant (2HPβCD treatment group: 33.5 ± 3.0%, and *minar2^fs139^* mutant: 58.0 ± 3.9% of those of controls), likely because that the cholesterol levels were also lower in the 2HPβCD treatment group (2HPβCD treatment group: 16.5 ± 2.4%, and *minar2^fs139^* mutant: 53.9 ± 5.2% of those of controls). When the *minar2^fs139^* mutants were treated with 2HPβCD, the cholesterol levels in the hair bundles and the AM1-43 labeling intensities were further reduced to levels in the 2HPβCD-treated wild type controls (***Figure 5A-B***). These data indicated that decreasing cholesterol levels in hair cells by 2HPβCD mimicked the defects in the *minar2^fs139^* mutant. In addition, 2HPβCD treatment aggravated hair cell defects in the *minar2^fs139^* mutant.

**Figure 5.**
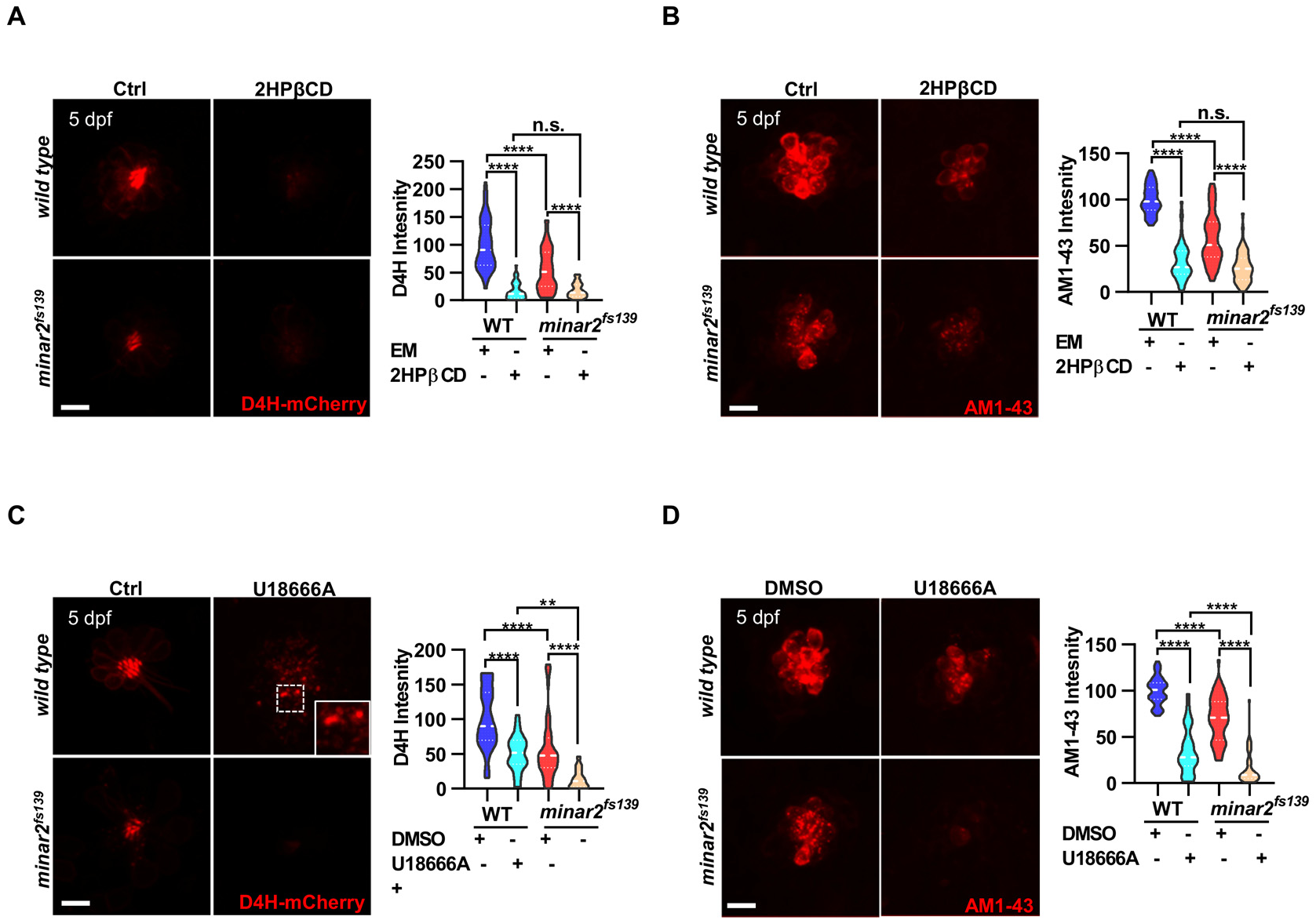
Lowering cholesterol levels aggravates hair cell defects in *minar2* mutant. **(A)** Quantification of D4H-mCherry intensity in wild type and mutant *minar2^fs139^* larvae after 2HPβCD treatment. The lateral line L3 neuromast were imaged and quantified. n = 45, 41, 52, and 51. F(3, 110.2) = 74.76, p<0.0001. **(B)** Quantification of AM1-43 labeling in wild type and mutant *minar2^fs139^* larvae after 2HPβCD treatment. n = 49, 46, 49, and 51. F(3, 155.4) = 124.4, p<0.0001. **(C)** Quantification of D4H-mCherry intensity in wild type and mutant *minar2^fs139^* larvae after U18666A treatment. n = 40, 31, 42, and 33. F(3, 111.3) = 39.34, p<0.0001. **(D)** Quantification of AM1-43 labeling in wild type and mutant *minar2^fs139^* larvae after U18666A treatment. n = 53, 50, 52, and 46. F(3, 169.9) = 158.0, p<0.0001. EM: embryonic medium, solvent control groups for 2HPβCD treatment (A and B); DMSO: solvent control groups for 2HPβCD treatment U18666A treatment (C and D); Multiple comparison significance values are indicated on the graph. Scale bars represent 10 μm. ***Figure 5-source data 1*** **Effects of decreasing cholesterol levels on hair cells in *minar2* mutant** Figure 5A-5D. Figure 5-source data 1.xlsx

We next used U18666A, a cationic amphiphile that inhibits Niemann-Pick C1 (NPC1) protein and cholesterol egress from lysosomes (Lu et al., 2015). Unlike a general reduction of cholesterol levels after 2HPβCD treatment, treatment with U18666A caused strong accumulation of cholesterol in the lysosome lumen (***Figure 2-figure supplement 1F***). Nevertheless, the D4H-mCherry labeling intensities were significantly decreased after the U18666A treatment (50.2 ± 4.6% of those of control, ***Figure 5C***). This was due to most cholesterol molecules were trapped inside lysosome and no longer accessible for D4H-mCherry, which was located in the cytoplasm. Treatment with U18666A also altered the distribution of the D4H-mCherry signals and gave rise to D4H-mCherry labeled endo-membrane particles at basolateral regions in the hair cells (***Figure 5C***, figure insert). Similar to the 2HPβCD treatment group, U18666A treatment strongly decreased the intensities of AM1-43 labeling, to levels similar in the 2HPβCD treatment group (***Figure 5D***). When the *minar2^fs139^* mutant were treated with U18666A, the cholesterol levels and the AM1-43 labeling intensities were further reduced to significantly lower levels than those in the U18666A-treated wild type controls (***Figure 5C-D***). Thus, there was an additive effect between the U18666A treatment and the *minar2^fs139^* mutation.

### Increasing cholesterol levels rescue hair cell defects and hearing in *minar2* mutant

We then tested whether increasing cholesterol levels had any effects on the hair cells. We first attempted treating wild type larvae with a 2HPβCD/cholesterol complex and found the 2HPβCD/cholesterol complex precipitated out in medium, likely due to the incubation temperature of 28.5 ℃ for proper zebrafish development (data not shown). We next used efavirenz, a cholesterol 24S-hydroxylase (CYP46A1) inhibitor, which can increase intracellular cholesterol levels by inhibiting hydroxylation of cholesterol and subsequent removal of cholesterol from cell (Petrov & Pikuleva, 2019). We found efavirenz consistently increased cholesterol levels in the neuromast hair cells (***Figure 6A***). We observed that treating the wild type larvae with efavirenz caused small increases of D4H-mCherry signals (111.2 ± 11.2% of controls, p=0.807) and AM1-43 labeling (107.4 ± 3.6% of controls, p=0.400) in neuromast hair cells (***Figure 6***), likely because the cholesterol homeostasis mechanism was intact in the wild type. In contrast, after treatment with efavirenz there were clear increases of D4H-mCherry signals in the hair bundles (***Figure 6A***, from 55.8 ± 7.0% to 88.7 ± 8.4% of those of controls, p=0.021), increases of AM1-43 labeling in hair cells in the 5 dpf *minar2^fs139^* mutant (***Figure 6B***, from 61.8 ± 2.9% to 96.1 ± 3.1% of those of controls, p<0.0001), and both were restored to the levels observed in the wild type controls (p=0.269 and p=0.115). We also found the treatment with efavirenz rescued the abnormally enlarged vesicles seen in the 8 dpf *minar2^fs139^* mutant (***Figure 6C***). We further tested voriconazole, another CYP46A1 inhibitor, and observed that the decreased AM1-43 labeling (***Figure 6-figure supplement 1A***) and the abnormal distribution of D4H-mCherry (***Figure 6-figure supplement 1B***) were similarly reversed after the voriconazole treatments.

**Figure 6.**
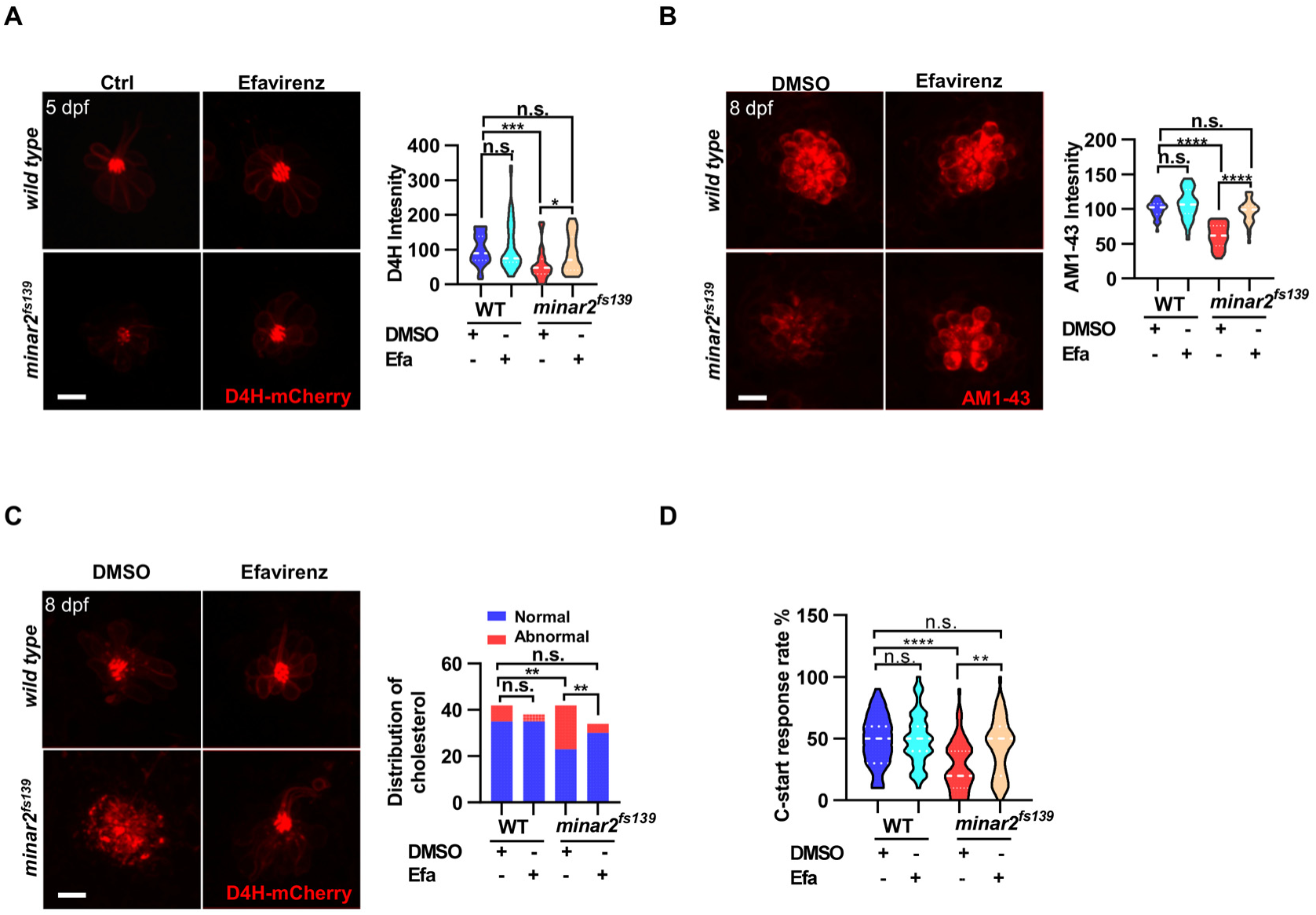
Increasing cholesterol levels rescue hair cell defects and hearing in *minar2* mutant. **(A)** Quantification of D4H-mCherry in hair cells of wild type and *minar2^fs139^* zebrafish after efavirenz treatment. The lateral line L3 neuromast were imaged and quantified. n = 40, 41, 42, and 41. F(3, 127.8) = 7.557, p<0.001. **(B)** Quantification of AM1-43 labeling in wild type and mutant *minar2^fs139^* larvae after efavirenz treatment. n = 31, 33, 33, and 30. F(3, 115.7) = 46.65, p<0.0001. **(C)** Effects of efavirenz treatment on appearance of abnormally enlarged vesicles in hair cells of wild type and mutant *minar2^fs139^* zebrafish. n = 42, 38, 42, and 34. χ^2^=20.92, df = 3, p<0.001. **(D)** Effects of efavirenz treatment on C-start response rates for wild type and *minar2^fs139^* mutants (n = 48 for all 4 groups. p < 0.0001, Kruskal-Wallis test). DMSO: solvent control groups; Efa: efavirenz treatment groups. Multiple comparison significance values are indicated on the graph. Scale bars represent 10 μm. ***Figure 6-source data 1*** **Effects of increasing cholesterol levels on hair cells in *minar2* mutant** Figure 6A-6D; Figure 6-figure supplement 1A-1B. Figure 6-source data 1.xlsx

We then examined whether efavirenz treatment rescued hearing defects in the *minar2^fs139^* mutant. We found that the compromised short-latency C-start (SLC) response rates in the *minar2^fs139^* mutant were rescued to levels in the wild type controls with the efavirenz treatment (***Figure 6D***). Thus, both the mechanotransduction (AM1-43 staining) and the hearing were restored after the decreased cholesterol levels were restored by the treatment with efavirenz.

### Minar2 interacts with cholesterol in vitro

The pharmacological treatment studies above strongly suggested Minar2 may function through the regulations of cholesterol. We next examined whether Minar2 interacted with cholesterol directly. The CSD of caveolin binds cholesterol through a so-called CRAC motif (Epand et al., 2005; Le Lan et al., 2010; Liu et al., 2016). We identified three CRAC (and its analog CARC) motifs in the primary sequences of Minar2 orthologs (***Figure 7A*** and ***Figure 1-figure supplement 1A***). We reasoned that if Minar2 interacted with cholesterol directly with the CRAC/CARC motif, point mutations of the critical aromatic residues (Y/F/W) within these motifs to alanine residues (Fantini et al., 2016; Li & Papadopoulos, 1998; Paschkowsky et al., 2018) should compromise association between Minar2 and cholesterol. We used the HEK293 cell to test this hypothesis in vitro, as GFP-Minar2 expressed in the cell strongly recruited cholesterol to perinuclear region and also increased intracellular cholesterol labeling (***Figure 3B***). We transfected HEK293 cells with wild type (MINAR2^1-190^) and point mutation construct (MINAR2^YW-A^), then stained the cholesterol with filipin (***Figure 7B***). We observed that the point mutation of MINAR2^YW-A^ compromised the recruitment of cholesterol to the perinuclear region (decreased from 24.58 ± 1.30 au in MINAR2^1-190^ group to 14.85 ± 0.90 au in MINAR2^YW-A^ group, p<0.0001) and the intensity of intracellular cholesterol labeling (from 103.9 ± 4.86 au in MINAR2^1-190^ group to 86.43 ± 3.21 au in MINAR2^YW-A^ group, p<0.01). These effects were not due to changes in expression levels of the point mutation construct, since it was expressed at similar level as the wild type construct (***Figure 7C***).

**Figure 7.**
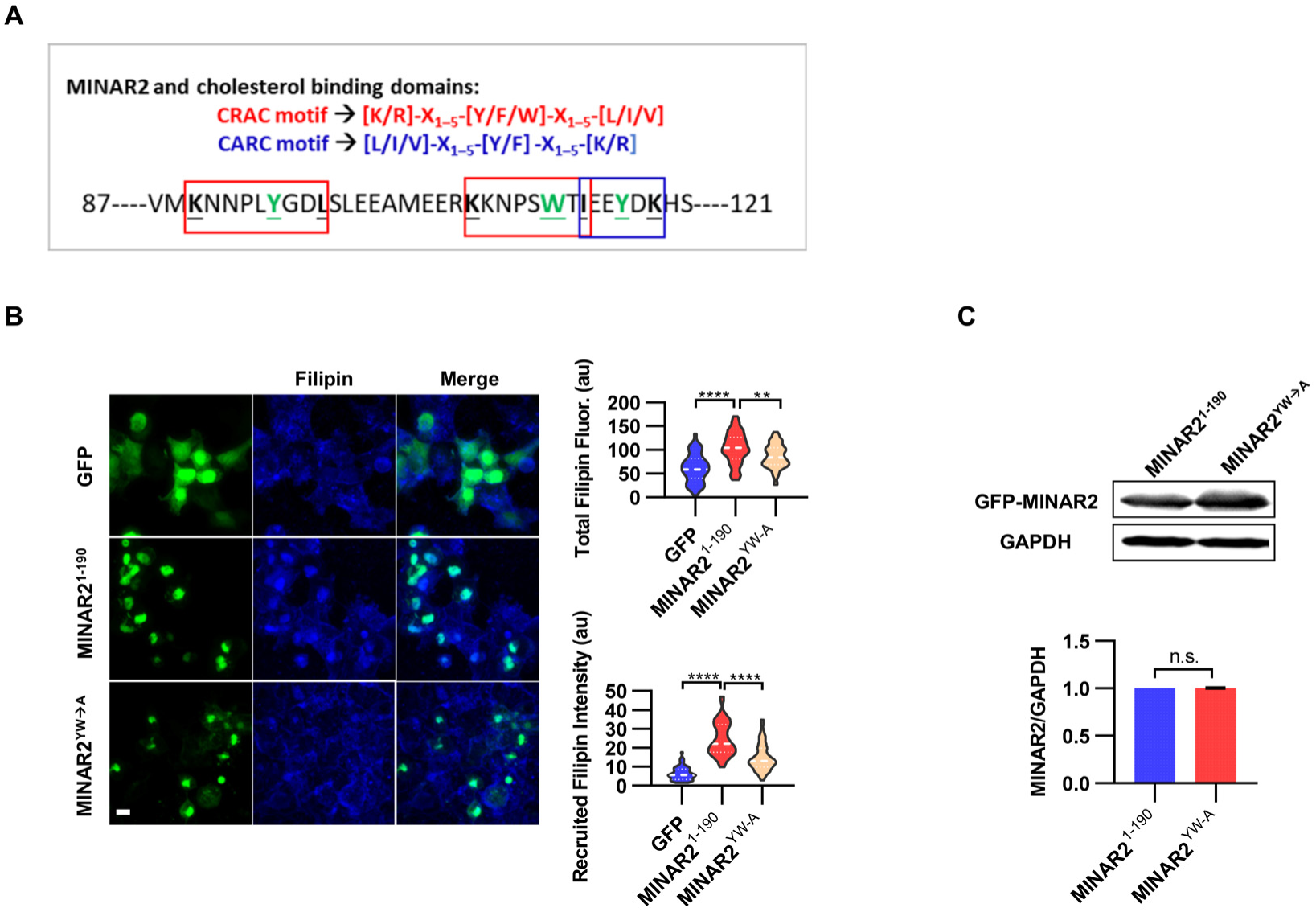
Minar2 interacts with cholesterol in vitro. **(A)** Cholesterol recognition motifs in Minar2 primary sequence. The sequences for cholesterol-recognizing amino-acid consensus (CRAC, red letter and box) and its analog CARC (blue letter and box) are indicated. The critical aromatic resides (Y/W, in green letters) were mutated to alanine in the point mutation MINAR2^YW-A^ construct. **(B)** Effects of critical aromatic reside mutation on the levels and distributions of filipin labeling in cultured cells. HEK293 cells were transfected with GFP alone (GFP), full length (GFP-MINAR2^1-190^), or the point mutation construct (GFP-MINAR2^YW-A^). n = 44, 46 and 62, for total filipin fluorescence, F(2, 149) = 23.80, p<0.0001; for recruited filipin intensity, F(2, 105.5) = 79.28, p<0.0001. Multiple comparison significance values are indicated on the graph. Scale bars represent 10 μm. **(C)** Immunoblot analysis of the expression levels of the full length construct (GFP-MINAR2^1-190^) and the point mutation construct (GFP-MINAR2^YW-A^) in HEK293 cells. Expression levels relatively to GAPDH were quantified. t=0.7498, df=2, p=0.532. ***Figure 7-source data 1*** **Interaction between Minar2 and cholesterol** Figure 7B-7C. Figure 7-source data 1.zip

In addition, the recently released MINAR2 structure model from AlphaFold2 (Tunyasuvunakool et al., 2021) allowed us to carry out computational docking to study probable Minar2-cholesterol interactions. We found the docking site with the best binding free energy (ࢤ7.1 kJ/mol) for cholesterol was indeed located in the conserved sequence pattern that matched to the CSD of caveolin, and the mutated aromatic residues (W112 and Y117) above were involved in the interaction between Minar2 and cholesterol (***Figure 7-figure supplement 1***).

## Discussion

We provide evidence that Kiaa1024L/Minar2 is required for normal hearing in the zebrafish. Our results suggest that Kiaa1024L/Minar2 protein, located on the apical endo-membranes and stereocilia membranes, is responsible for apical distribution of cholesterol to the hair bundle, and this apical distribution of cholesterol is essential for the normal morphogenesis and function of the hair bundle. Consistent with this model, loss of *minar2* markedly reduces cholesterol distribution in the hair-bundles, causes longer and thinner hair bundles, and gives rise to enlarged and aggregated apical lysosomes. These defects in *minar2* mutant likely compromise mechanotransduction, reduce the number of hair cells, and result in hearing loss. Our model is further supported by results from pharmaological interventions. The effects of *minar2* loss-of-function is mimicked by drug treatment that reduces the cholesterol levels in the hair bundles, while these effects are reversed by treatment increasing the cholesterl levels.

*Kiaa1024L/Minar2* is a previously understudied gene and present in the vertebrate species only. Previous mouse genetic screen for hearing loss gene showed knockout of *Kiaa1024L/Minar2* causes profound deafness (Bowl et al., 2017; Ingham et al., 2019). Our study presented here, together with the results in the mouse model, highlight that *Kiaa1024L/Minar2* is essential for normal hearing in vertebrates. Our results with mutant zebrafish reveal that the hearing defects exist at the level of hair cells. This conclusion is consistent with the highly enriched expression of *Kiaa1024L/Minar2* in the hair cells in the vertebrate inner ears, shown by our in situ hybridization results and our surveys of FACS and single cell RNA sequencing data (Barta et al., 2018; Elkon et al., 2015; Erickson & Nicolson, 2015; Liu et al., 2018; Steinhart et al., 2022). Outside the inner ear, *Kiaa1024L/Minar2* is also expressed in various regions in the brain tissue, and a previous study showed loss of *Minar2* in mice impairs motor function and reduces the number of tyrosine hydroxylase-positive neurons (Ho et al., 2020).

The degree of hearing loss in mutant *minar2^fs139^* zebrafish larvae is not as severe as it was in the adult knockout mouse. Since the hearing function of young *Minar2* knockout mice was not reported, it is not known whether the profound deafness in mice is progressive or adult-onset. In mutant *minar2^fs139^* zebrafish, the reduction of inner ear hair cell numbers is progressive, worsening from no changes in larval stage to 30% reduction in adult stage. We note that unlike the mouse model, in which the hair cells are generated in early development and cannot regenerate in adult stage (Atkinson et al., 2015; Basch et al., 2016; Driver & Kelley, 2020), in zebrafish the hair cells are continuously generated until late in adulthood (Wang et al., 2015). Investigation in the mouse model is required to determine whether there also is a progressive loss of hair cells and if the degree of loss is more extensive to account for the severe hearing loss in the mouse.

Defects in mechanotransduction, longer and thinner hair bundles, and enlarged apical lysosomes observed in *kiaa1024L/minar2* mutant all indicate that *Kiaa1024L/Minar2* acts primarily in the apical region and in the hair bundle in hair cells. GFP-tagged Minar2 protein is primarily localized to the the apical endo-membranes and stereocilia membranes, which are the appropriate locations to exert above functions. Previous studies showed that the apical region of the hair cell is crowded with endocytotic vesicles, including abundant lysosomes (Krey et al., 2018; Revelo et al., 2014; Spicer et al., 1999; Wiwatpanit et al., 2018) These endocytotic membrane bound structures are thought to transport ions and other materials to and fro towards the apical entry of the hair cells (Spicer et al., 1999), but the functions of these structures are not well characterized. One recent study showed that loss of lysosomal mucolipins in cochlear hair cells gives rise to abnormally enlarged lysosomes at the apical region. These aberrant lysosomes are toxic to cell and cause hair cell loss (Wiwatpanit et al., 2018).

At the apical end of the inner ear hair cell, the distribution pattern of the cholesterol probe D4H-mCherry in the hair bundle is striking. The D4H-mCherry probe is almost exclusively localized in the hair bundle; while the plasma membrane probes PM-GFP is localized both to basolateral and apical membranes. Previous study showed that stereocilia membrane is densely covered with filipin-induced deformation in freeze facture preparation of hair cells, indicating presence of abundant cholesterol in the stereocilia membrane (Forge et al., 1988; Forge & Richardson, 1993). The exact function of this striking distribution pattern of cholesterol requires further investigation. Cholesterol is a major component of biological membranes, and it plays both structural and functional roles in the bio-membranes (Maxfield & van Meer, 2010; Subczynski et al., 2017). Previous studies showed that cholesterol may stiffen biological membranes (Chakraborty et al., 2020; Dimova, 2014). When cholesterol was depleted from hair cell membranes following methyl β cyclodextrin treatment, an apparent loss of structural integrity and “floppy” hair bundles were observed (Purcell et al., 2011). The biophysical properties of membranes may be more directly involved in the mechanotransduction. It has been suggested that the stereocilia tip membranes can serve as a component of the gating spring of the MET channel (Powers et al., 2014; Powers et al., 2012), although this has currently been debated (Bartsch et al., 2019). On the other hand, it is clear that cholesterol has direct or indirect effects on membrane proteins such as ion channels and G-protein coupled receptors (Harris, 2010). Previous studies showed that the mechanosensitive ion channel Piezo1 interacts with cholesterol and this interaction is essential for the spatiotemporal activity of Piezo1 (Buyan et al., 2020; Ridone et al., 2020).

Our model suggests that Kiaa1024L/Minar2 regulates the distribution and homeostasis of cholesterol to ensure normal hearing. Several lines of evidence indicate Kiaa1024L/Minar2 interacts with cholesterol: (1) the highly conserved sequence pattern found in Kiaa1024L/Minar2 orthologs matches the cholesterol binding CSD domain of caveolins; (2) Kiaa1024L/Minar2 recruits cholesterol in vitro, and point mutations in the predicted cholesterol binding sites abolish the recuitment of cholesterol; (3) computational docking study with atomic model of Kiaa1024L/Minar2 from AlphaFold2 shows reasonable binding free energy between cholesterol and the sequence pattern we identified. Neverthessless, further biochemical studies are required to demonstate direct binding between the Kiaa1024L/Minar2 protein and cholesterol.

It is currently unclear how Kiaa1024L/Minar2 regulates the distribution and homeostasis of cholesterol in hair cells. Kiaa1024L/Minar2 may be responsible or faciliate the apical transport of cholesterol to the hair bundles. The subcellular localization of Kiaa1024L/Minar2 in both the apical endo-membranes and the hair bundle membranes is consistent with a transporting role for Kiaa1024L/Minar2. The apical transport defects caused by loss of Kiaa1024L/Minar2 can explain the marked decreases of accessible cholesterol levels in the hair bundles. Because the accessible cholesterol is the critical determininant for the cholesterol homeostasis regulation (Lange & Steck, 2016; Lange et al., 2004), the decreases of accessible cholesterol is expected to bring about the up-regulation of genes downstream of Srebp2, which is indeed observed in the *minar2* mutant. Under this model, the observed enlarged aggreated lysosome and reduced number of hair cells are likeky secondary to the transportation defects. It will be important to examine the cholesterol dynamcis in vivo and to define the interacting proteins of Kiaa1024L/Minar2 to illustrate the detailed mechanism for the function of Kiaa1024L/Minar2.

Kiaa1024L/Minar2 protein belongs to a protein family that also includes the Kiaa1024/Minar1/Ubtor protein. This protein family is defined by a conserved but <u>u</u>ncharacterized <u>p</u>rotein <u>f</u>amily domain (UPF0258) located at the carboxy ends of the proteins. Our previous study showed that Kiaa1024/Minar1/Ubtor acts as a negative regulator of mTOR signaling through its interaction with the Deptor protein, an integral component of the mTORC complex (Zhang et al., 2018). Loss of *ubtor* gene in zebrafish causes motor hyperactivity and epilepsy-like behaviors by elevating neuronal activity and activating mTOR signaling (Wang et al., 2021). It has also been shown that *Ubtor/MINAR1* play roles in angiogenesis and breast cancer (Ho et al., 2018). Since the region in Kiaa1024/Minar1/Ubtor that interacts with the Deptor protein is missing in the Kiaa1024L/Minar2 protein, the regulation of mTOR signaling is likely a function specific to the Kiaa1024/Minar1/Ubtor protein. Importantly, a recent study showed that *Kiaa1024/Minar1/Ubtor* is a host factor crucial for the replication of pathogenic human coronaviruses, including the SARS-CoV-2 that causes the COVID-19 (Kratzel et al., 2021). Previous studies showed that the replications of positive-strand RNA viruses including hepatitis C virus (HCV) and SARS-CoV-2 require rearranged intracellular membrane structures called replication organelle (Twu et al., 2021). The SARS-CoV-2 replication organelle is composed of endoplasmic reticulum derived double membrane vesicles (DMV). Interestingly, we previously showed that Kiaa1024/Minar1/Ubtor protein is localized to the endoplasmic retiulum (Zhang et al., 2018). Furthermore, DMVs are enriched for cholesterol (Paul et al., 2013) and endosomal cholesterol homeostasis is important for the DMV formation and replication of various viruses (Glitscher & Hildt, 2021; Stoeck et al., 2018). Because the cholesterol-binding region that we identify in this study is conserved in the Kiaa1024/Minar1/Ubtor protein, it will be important to examine whether Kiaa1024/Minar1/Ubtor also regulates cholesterol distribution and homeostatsis to support SARS-CoV-2 replication.

In conclusion, our data provide evidence that cholesterol plays an essential role in the hair bundle, and Kiaa1024L/Minar2 regulates cholesterol distribution and homeostasis to ensure normal hearing. Because of the conserved requirement of *Kiaa1024L/Minar2* gene orthologs for hearing in both mouse and zebrafish, and that the human ortholog *MINAR2* is also spcifically expressed by the auditory hair hairs (Steinhart et al., 2022), it will be important to examine whether *KIAA1024L/MINAR2* gene mutations may underline hearing abnormality in human. Further studies of Kiaa1024L/Minar2’s regulation of cholesterol distribution and homeostasis in the zebrafish and mammalian model will provide new insights into the biological process for the morphogenesis and physiology of hair bundles and hearing loss. The involvement of cholesterol in the functions of the UPF0258 family proteins may also have implications in how *Kiaa1024/Minar1/Ubtor* regulates the replication of coronavirus, an urgent question of societal and scientific significance.

## Materials and Methods

### Key resources table

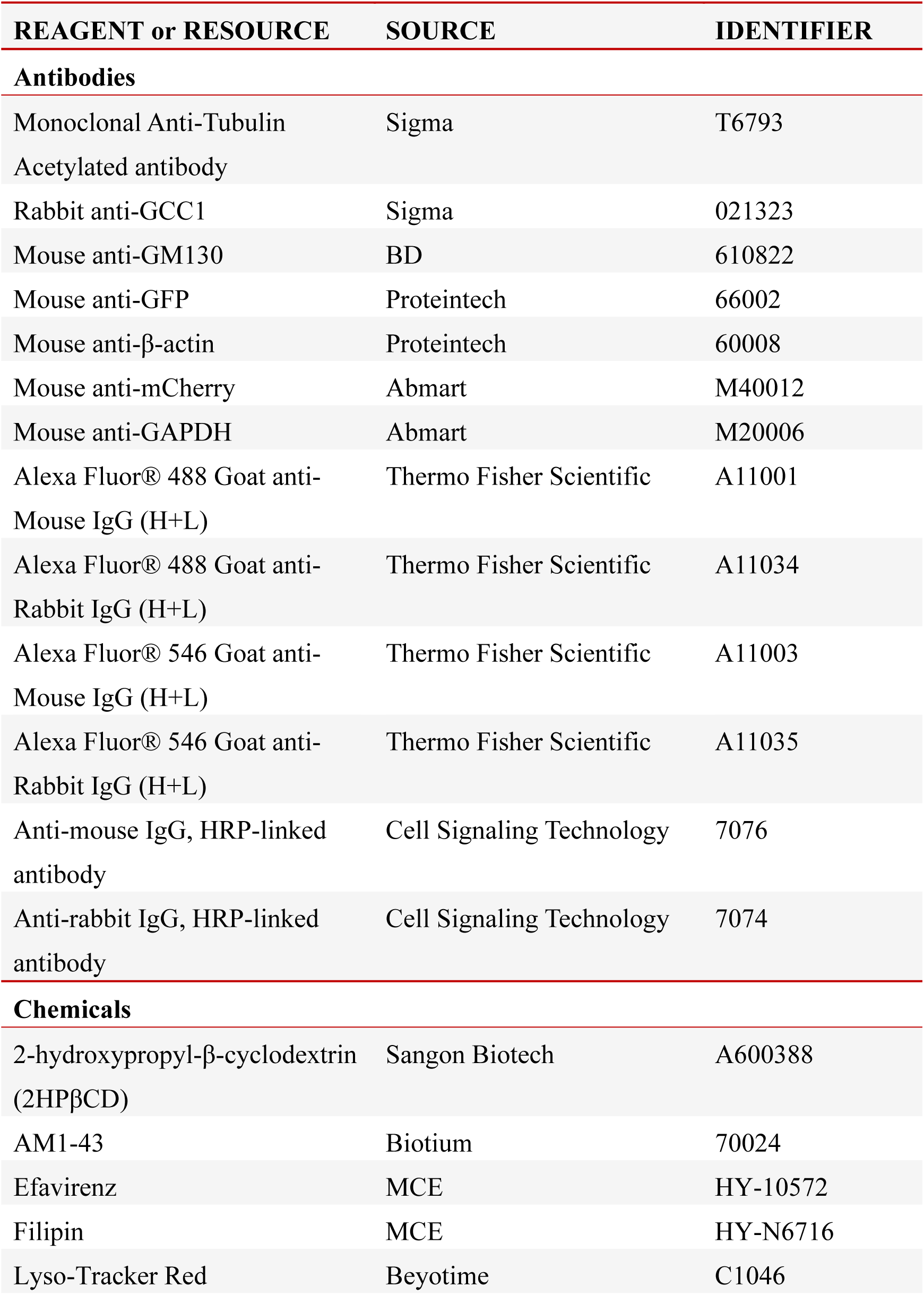
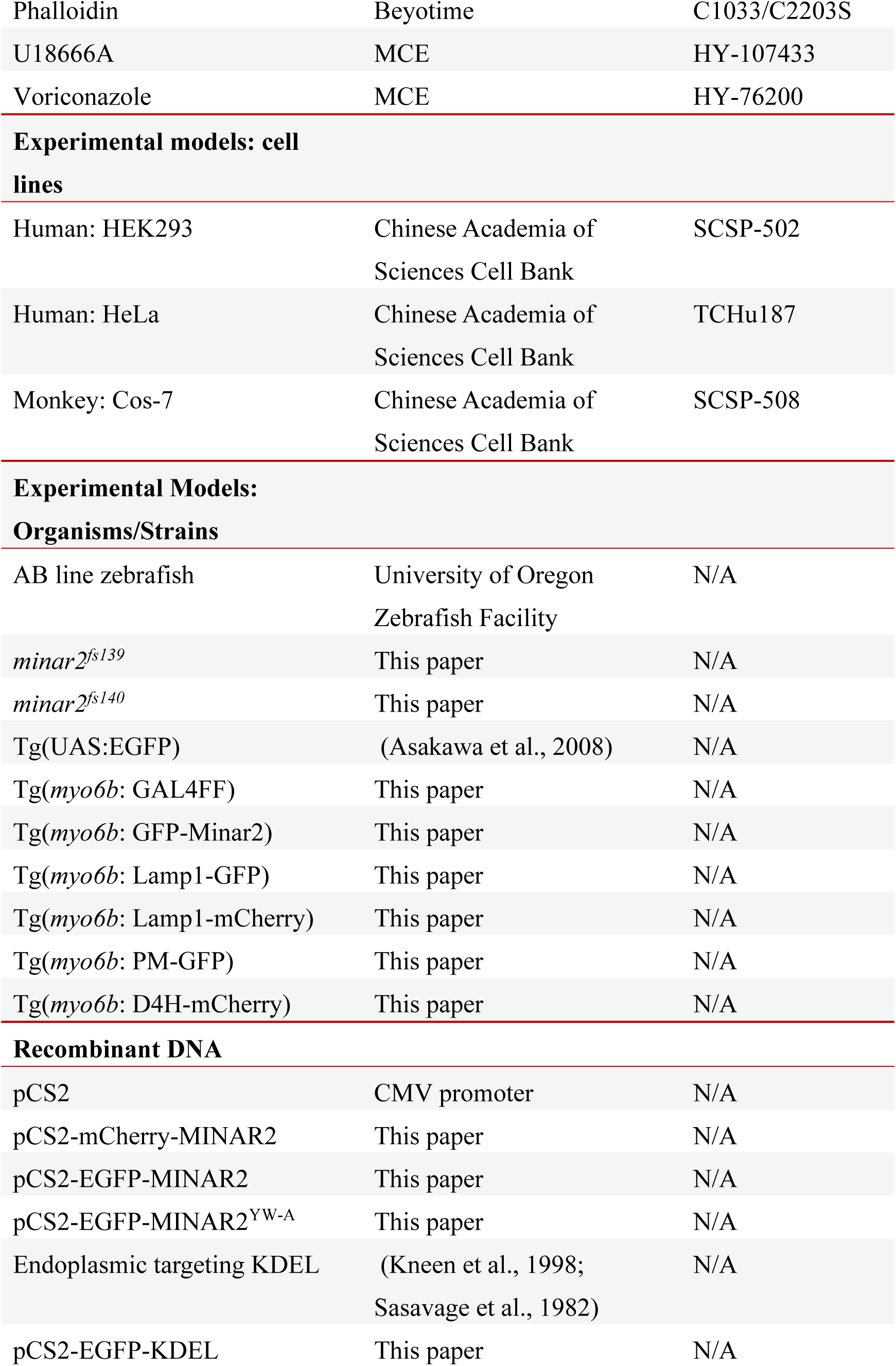
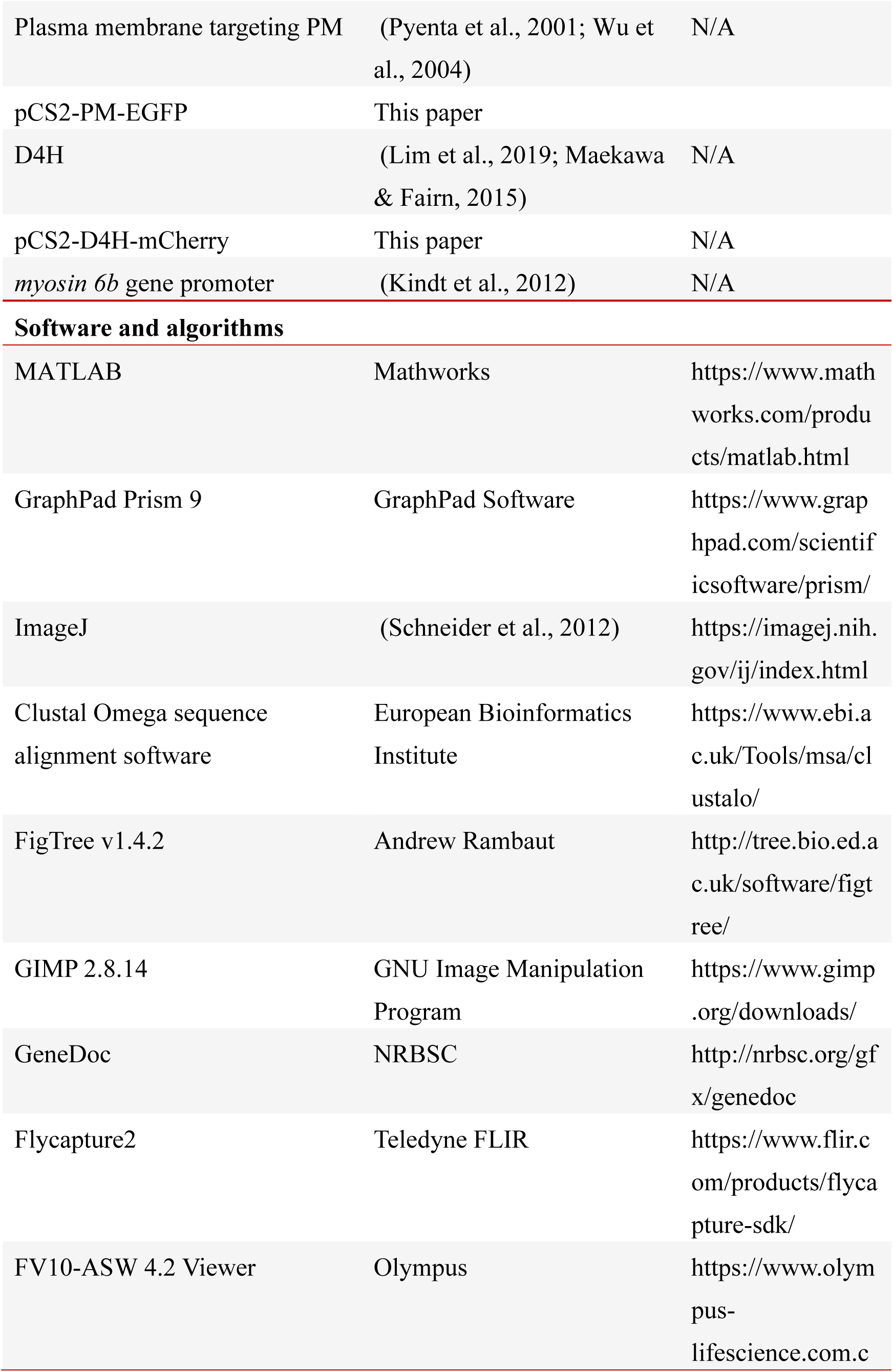
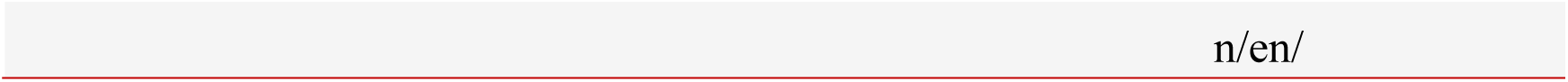

### Oligonucleotides

Primers for genotyping

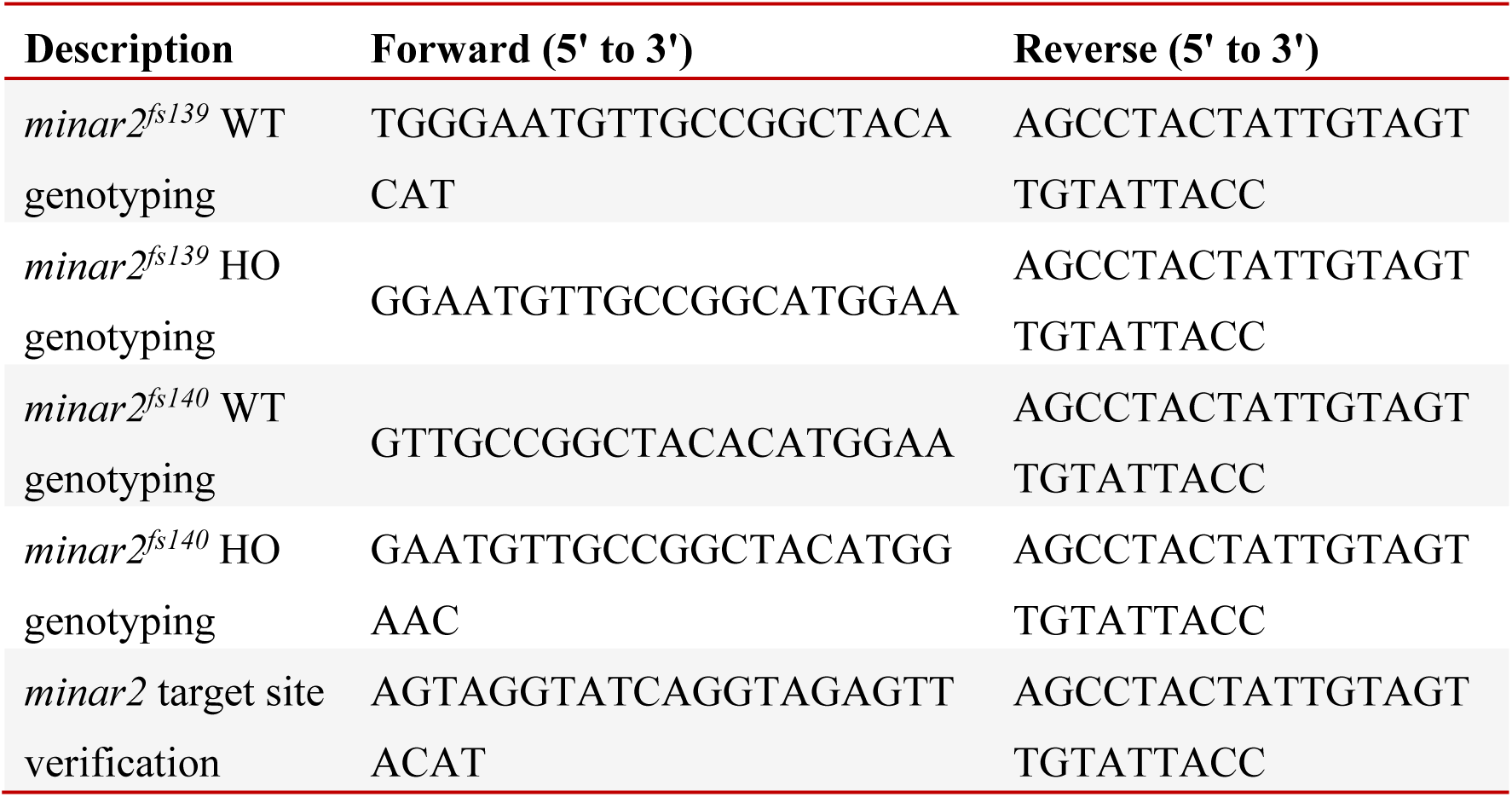

Primers for RT-PCR and probe

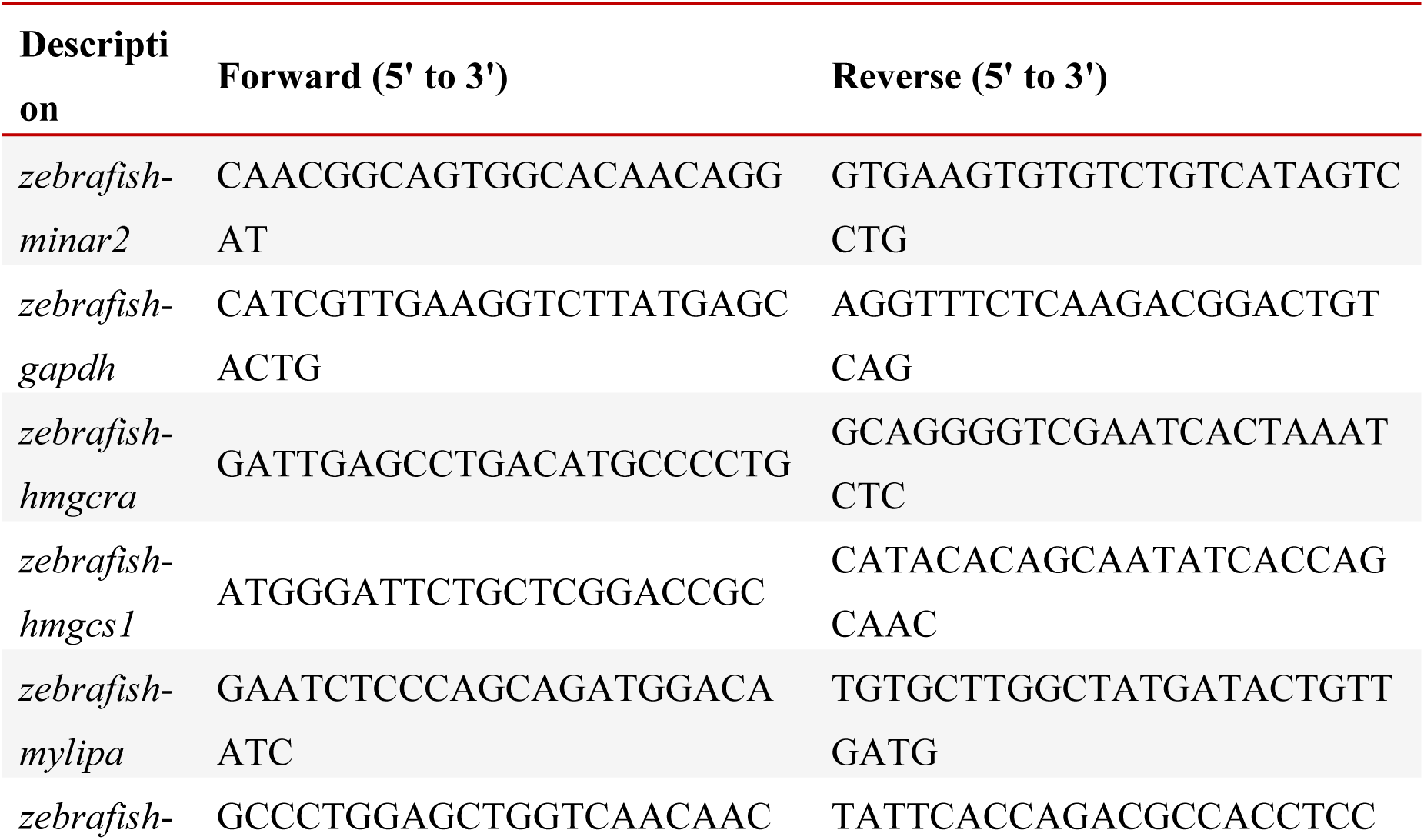
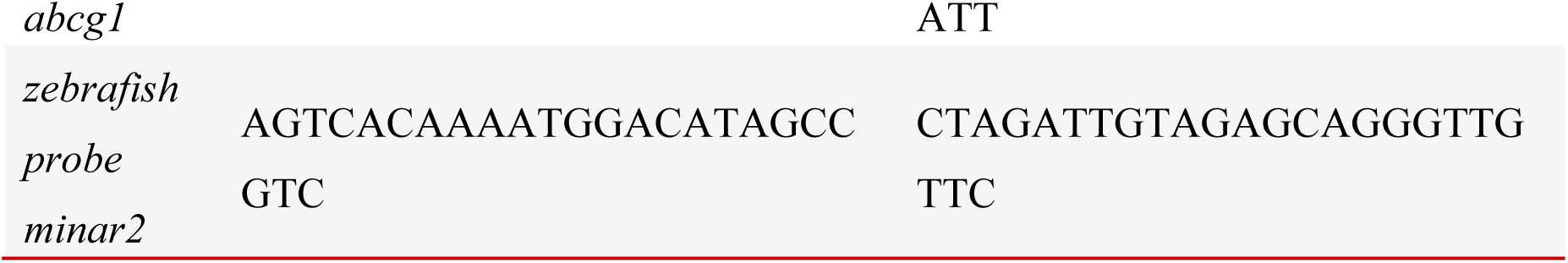

Primers for plasmid construction

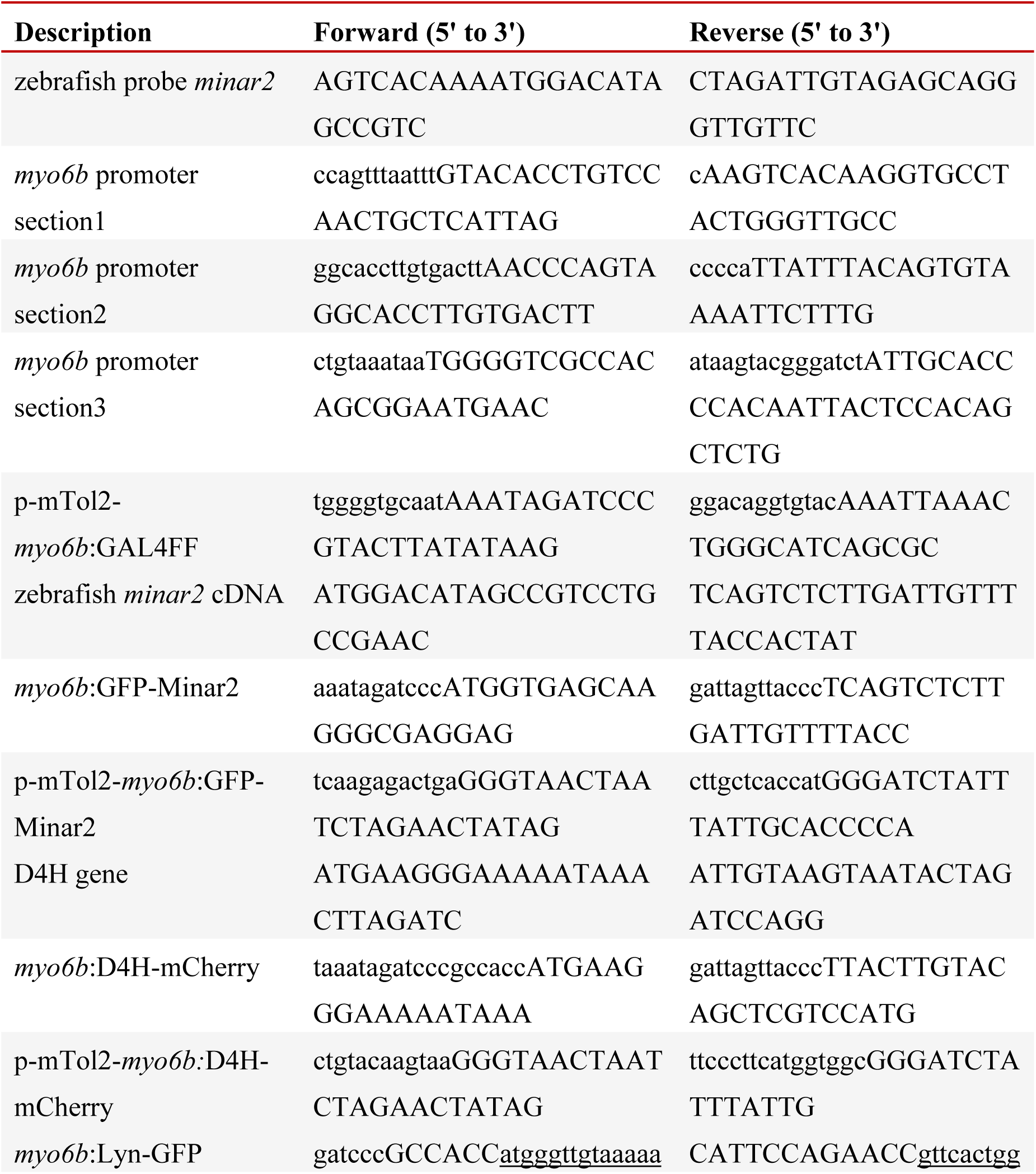
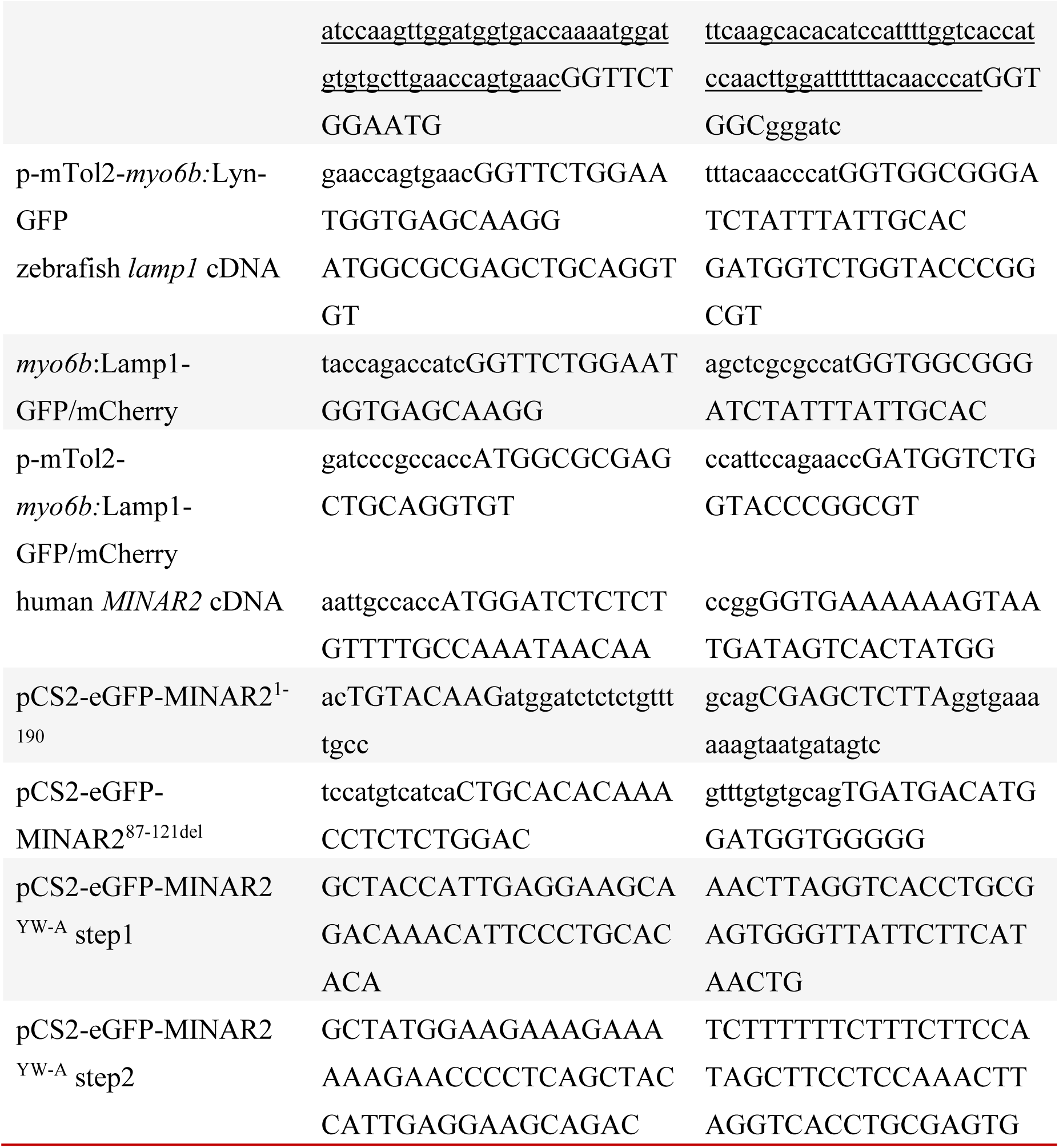

### Resources and materials availability

All constructs, transgenes, and reagents generated in this study are available from the Lead Contact without restriction. Further information and requests for resources and materials should be directed to and will be fulfilled by the Lead Contact, Gang Peng.

### Zebrafish strains

The animal use protocols were approved by the Fudan University Shanghai Medical College Institution Animal Care and Use Committee (130227-092, 150119-088 and 190221-147). All animals were handled in accordance with the Fudan University Regulations on Animal Experiments.

Zebrafish were maintained according to standard protocols (Westerfield, 2007). Lines used in this study included the AB, Tg(UAS:EGFP), Tg(myo6b:GAL4FF), Tg(myo6b:GFP-Minar2), Tg(myo6b:D4H-mCherry), and *minar2* mutant (*minar2^fs139^*and *minar2^fs140^*). The Tg(UAS:EGFP) line was a gift from Dr. Kawakami (Asakawa et al., 2008). Zebrafish larvae were obtained from natural spawning and fed paramecia beginning on 5 dpf. Larvae at 5-8 days post fertilization (dpf) of undifferentiated sex were examined.

### Cell lines

The HEK293, HeLa and COS-7 cell lines were obtained from the Cell Bank of the Chinese Academy of Sciences (Shanghai). HeLa cells were grown in MEM/EBSS (HyClone) medium. HEK293T and COS-7 cells were grown in DMEM/High glucose (HyClone) medium. The growth media were supplemented with 10% fetal bovine serum (FBS) (Biological Industries). For transfection of cells, Lipofectamine 3000 (Thermo Fisher Scientific) was used.

### Generation of *minar2* mutant

To disrupt the *minar2* gene in the zebrafish, targeted lesions were introduced into the zebrafish genome through CRISPR/Cas9 mediated gene modification. The sgRNA targeted sequence (5’-GGAATGTTGCCGGCTACACATGG-3’) was located in the first exon of *minar2* gene. Two *minar2* mutant lines (*minar2^fs139^*and *minar2^fs140^*) were each outcrossed with the AB line for 6 generations and then used in this study. Allele-specific primers (Oligonucleotide table) were used to genotype wild type and mutant siblings.

### Constructs and transgenic lines

Plasmid constructs were generated by standard molecular cloning with primers and oligonucleotides listed in Oligonucleotide table. Transgenic lines were generated using the Tol2 transposase-mediated method (Kawakami et al., 2000; Urasaki et al., 2006). Transgenic lines were outcrossed with the AB line for three generations before used in this study. For the Tg(myo6b:D4H-mCherry) line, the genomic insertion site of the transgene was mapped by high-efficiency thermal asymmetric interlaced PCR (Liu & Chen, 2007; Liu & Whittier, 1995) as previously described (Zhang et al., 2018).

### C-start response

C-start response was carried out as previously described (Zhang et al., 2018). Briefly, each well of a 4 by 4 test grid was filled with an individual larva of 5 or 8 dpf and 150 μL embryo medium. The larvae were allowed to adapt to the wells for 5 min prior to the start of the test. The microcontroller delivered 10 acoustic stimuli (200 Hz, 50 ms), with randomized inter-stimulus intervals (varied from 40 to 120 seconds) to minimize the adaptation of the zebrafish. The video sequence was inspected by person blind to the genotype condition, and C-start responses were recorded if the zebrafish performed the C-bend within 25 ms after the start of the stimulus. The C-start response rates were calculated by the numbers of C-start performed out of the 10 stimuli delivered.

### Auditory evoked potential (AEP) recording

AEP recordings were carried out following procedures from published studies with modifications (Higgs et al., 2002; Wang et al., 2015). Each zebrafish larva (7-8 dpf) was mounted dorsal-side up in 1% low melt point agarose at the center of a 35 mm glass-bottom petri dish, then covered with 1 ml of distilled water. A recording electrode was inserted into the dorsal surface of brainstem, between the two otic vesicles. A reference electrode was placed near the surface of the forebrain. Electrodes were insulated stainless steels (160 µm in diameter), and the dimensions of the exposed tip were approximately 80 µm in height, and 50 µm (base) to 5 µm (end) in diameter (Zhongyan, Beijing). AEP stimuli generation and signal acquisition were controlled by custom MATLAB scripts and a custom C++ program. AEP stimuli consisted of Blackman-windowed tone pips (10 ms in duration) of varying frequencies (100, 200, 400, 600, 800 and 1000 Hz) and were presented in descending sound levels in 2-5 dB decrements. AEP potentials were amplified 80,000x (TengJun Amp, Xi’an), digitized (Art Technology, Xi’an), then processed and averaged by custom MATLAB scripts.

### Mechanotransduction assay with AM1-43 labeling

Larvae were incubated in 0.2 μM AM1-43 (Biotium, California) in low calcium Ringer’s solution (LCR) [140 NaCl, 2 KCl, 0.1 CaCl_2_, 5 D-glucose, and 5 HEPES (mM)] at room temperature for 3 min and then washed in LCR for 2 x 5 min. The samples were then fixed in 4% paraformaldehyde for 1 hr at room temperature before storage and imaging. The lateral line neuromast L3 was imaged and the voxels within a bounding box (30 x 30 x 15 µm in 10 optical sections) containing the hair cells were processed for quantification using custom MATLAB scripts.

### Phalloidin staining

Larvae were fixed in 4% paraformaldehyde/1x PBS for 1 hr at room temperature then overnight at 4 ℃. Fixed specimens were permeabilized in 1% Triton-X100/1x PBS for 1 hr at room temperature, and then incubated with fluorescence-conjugated phalloidin (Beyotime, Shanghai, 1:200 in 1% Triton-X100/1x PBS) for 2 hr at room temperature. DAPI was used to counter-stain the nuclei.

### Lyso-Tracker Red staining

To stain cultured cells, cells were grown in glass-bottom petri dishes. Lyso-Track Red (Beyotime, Shanghai) was diluted (1:2000) in cell culture medium and preheated to 37 ℃ before replacing medium in the glass-bottom petri dishes. Cells were incubated in medium containing Lyso-Track Red for 10 min at 37 ℃, and then washed with culture medium before imaging. To stain larvae, Lyso-Tracker Red was diluted (1:2000) in the embryonic medium (EM). Larvae were incubated in EM containing Lyso-Tracker Red for 1.5 hr at 28 °C. After the staining, larvae were rinsed three times with EM before imaging. All staining steps were performed in the dark.

### Filipin staining

Filipin (MCE, Shanghai) stock was prepared in DMSO (50 mg/ml). To stain cultured cells, cells grown on coverslips were fixed with 4% formaldehyde/1x PBS for 10min at room temperature. Fixed cells were stained with freshly prepared filipin working solution (0.25 mg/ml in 1xPBS) for 15 min at room temperature. To stain larvae, larvae were fixed in 4% paraformaldehyde/1x PBS for 2 hr at room temperature then overnight at 4 ℃. Fixed specimens were incubated with filipin working solution for 2 hr at room temperature. Samples stained with filipin were imaged with a 405 nm laser. Quantification of filipin staining was carried out with custom MATLAB scripts.

### Immunofluorescent staining

Larvae were fixed in 4% paraformaldehyde/1x PBS for 1 hr at room temperature then overnight at 4 ℃. Fixed samples were treated with methanol overnight at -20 ℃, and then step-washed in PBST (1x PBST/0.1% Tween-20). Fixed larvae were permeabilized in 1% Triton X-100/1x PBS for 1 hr at room temperature. Cultured cells grown on coverslips were fixed with 4% paraformaldehyde/1x PBS for 10 min at room temperature, and then post-fixed with methanol at -20 ℃ for 15 min. Fixed cells were permeabilized with 0.5% Triton X-100/1x PBS for 7 min. Fixed and permeabilized samples were blocked in PBS containing 2% sheep serum, 2% goat serum, 0.2% BSA, and 0.1% Tween-20. Blocked samples were washed, incubated with primary antibodies, and then with secondary antibodies.

### Quantification of fluorescence signal

Fluorescence signals were quantified with custom MATLAB scripts. In brief, raw image data were imported using Bio-Formats (Linkert et al., 2010), segmented with an adaptive thresholding method, then the selected pixels were quantified.

### Adult inner ear dissection and analysis

Dissections of inner ears in adult zebrafish were carried out following published protocols (Liang & Burgess, 2009; Monroe et al., 2016). The dissected inner ear sensory epithelia (utricle and saccule) were incubated with fluorescence-conjugated phalloidin (Beyotime, Shanghai) for 30 min at room temperature. For quantification, three images were acquired for each utricle (area 1: center of the extrastriolar region; area 2 and 3: lateral and medial striolar region, respectively), or for each saccule (area1, 2, and 3: 5%, 50% and 90% positons along the anterior-posterior axis). Hair bundles were counted within a 50 x 50 μm region centered on each image using ImageJ multi-point tool.

### Immunoblot analysis

Cultured cells or zebrafish larvae were lysed in 2 x SDS sample buffer, and then boiled at 95 ℃ for 10 min to extract proteins. Extracted proteins were resolved by 6-20% SDS-PAGE, and then tank-transferred onto nitrocellulose membranes. Blotted membranes were probed with primary antibodies overnight and detected with HRP conjugated secondary antibodies.

### Pharmacological treatment

2-hydroxypropyl-β-cyclodextrin (2HPβCD) stock (200 mM) was prepared in EM. Larvae were treated with 7.5 mM 2HPβCD diluted in EM from 2 dpf to designated time, and the treatment media were renewed every day. U18666A stock (10 mM) was prepared in DMSO. Larvae were treated with 7 µM U18666A diluted in EM from 2 dpf to designated time, and the treatment media were renewed every day. Efavirenz stock (20 mM) and voriconazole stock (10 mg/ml) were prepared in DMSO. Larvae were treated with 2 µM efavirenz, or 1 μg/ml voriconazole diluted in EM from 2 dpf to designated time. For solvent controls, larvae were exposed to the same concentrations of EM or DMSO as in the drug treated groups.

### AutoDock analysis of cholesterol interaction

Docking was performed using AutoDock 4.2 and AutoDockTools (ADT 1.5.7) as previously described (Forli et al., 2016; Morris et al., 2009). The PDB file for MINAR2 was downloaded from Alphafold (https://alphafold.com/entry/P59773). The cholesterol ligand was prepared by importing experimental model coordinates for cholesterol from the Ligand-Expo of RCSB (http://ligand-expo.rcsb.org/reports/C/CLR/CLR_model.pdb).

### Statistical analysis

Sample sizes were estimated based on data variances and effect sizes in pilot experiments. Data analyses and plots were carried out using GraphPad Prism 9. Significance values are denoted as, *: *p*<0.05, **: *p*<0.01, ***: *p*<0.001, ****: *p*<0.0001. All quantified data are from at least three biological repeats. Sample sizes for each figure are given in the figure legends. Significance of differences was assessed by two-tailed Student’s *t*-test, one sample *t*-test, one way ANOVA (ordinary or Brown-Forsythe), two way ANOVA, or non-parametric analysis when appropriate.

## Acknowledgments

We thank Dr. Teresa Nicolson for providing the sequence information of the *myo6b* constructs. We thank Drs. Wei-Jun Pan, Qing-jian Han, and Ying Cao, and members of the laboratory for helpful discussions. We thank X Li and C Han for fish care. This work was supported by National Key Research and Development Program of China (2018YFA0801000), National Natural Science Foundation of China (31571067), Shanghai Municipal Science and Technology Major Project (No.2018SHZDZX01), ZJ Lab, and Shanghai Center for Brain Science and Brain-Inspired Technology.

---

**Figure 1-figure supplement 1.**
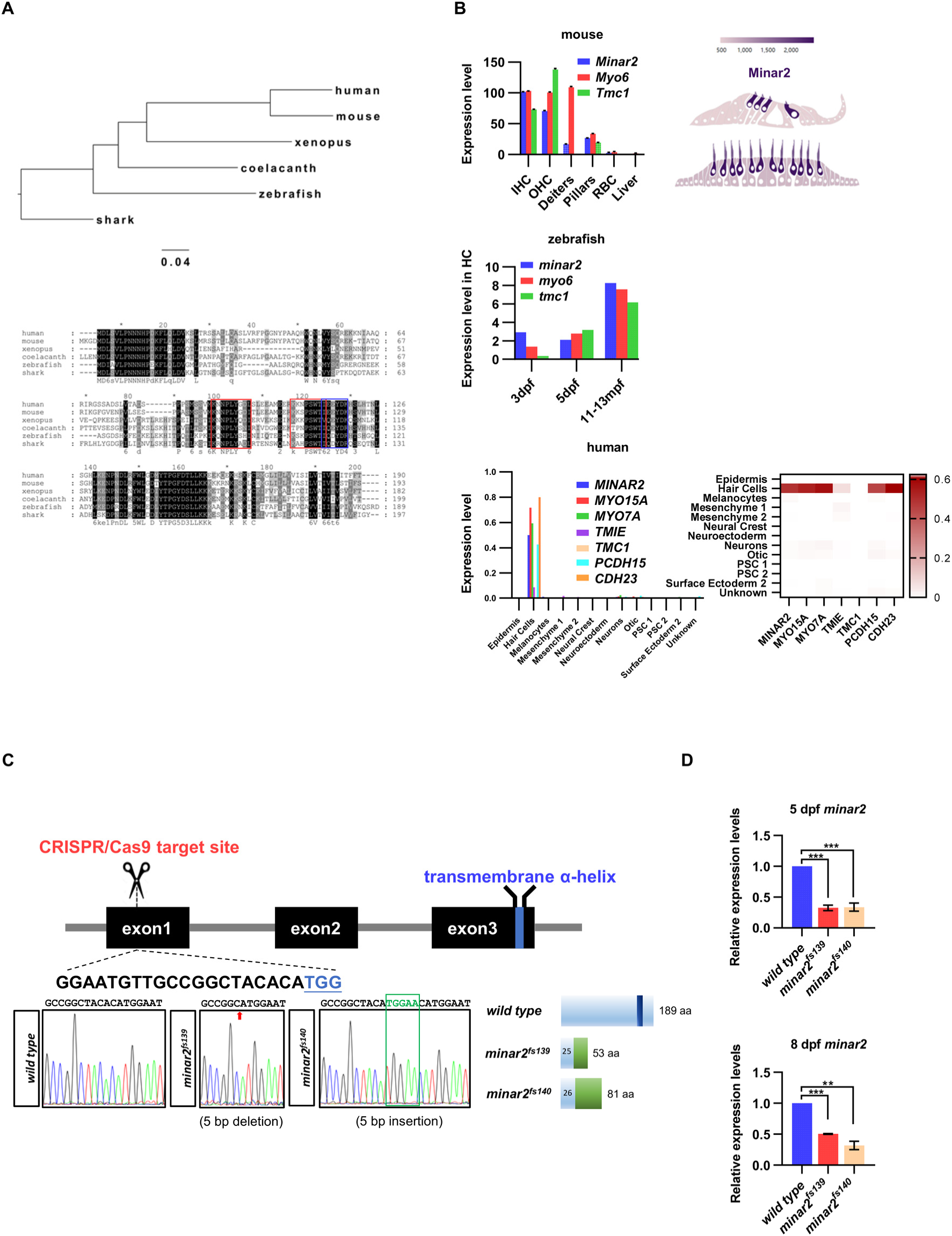
Expressions of *minar2* orthologs in hair cells and generation of *minar2* mutant alleles in the zebrafish. **(A)** Phylogenetic tree and multiple sequence alignment of *minar2* orthologs. No *minar2* homologs were found outside the vertebrates. The sequences matched to the cholesterol-recognizing amino-acid consensus motif (CRAC, red box) and its analog CARC (blue box) are indicated. **(B)** *minar2* orthologs are highly expressed in the hair cells. Expression data were taken from published sources: mouse (Elkon et al., 2015; Liu et al., 2018), zebrafish (Barta et al., 2018; Elkon et al., 2015; Erickson & Nicolson, 2015), and human inner ear organoid (Steinhart et al., 2022). **(C)** Targeted disruption of the zebrafish *minar2* gene by CRISPR/Cas9. The sequence for gRNA target site is indicated under exon1 box. Sequencing chromatographs are marked to show altered nucleotides in the *minar2* loci. Schematic representations of wild-type and mutant Minar2 proteins are shown next to the sequencing chromatographs. **(D)** Real-Time quantitative reverse transcription PCR analysis of *minar2* expression levels. Expression levels relative to β-actin levels were normalized to the wild type control group. For 5 dpf samples, wild type versus *minar2^f139^*: t=14.94, df=4, ***p<0.001; wild type versus *minar2*^f140^: t=10.29, df=4, ***p<0.001. For 8 dpf samples, wild type versus *minar2^f139^*: t=75.38, df=2, ***p<0.001; wild type versus *minar2*^f140^: t=10.16, df=2, **p<0.01.

**Figure 1-figure supplement 2.**
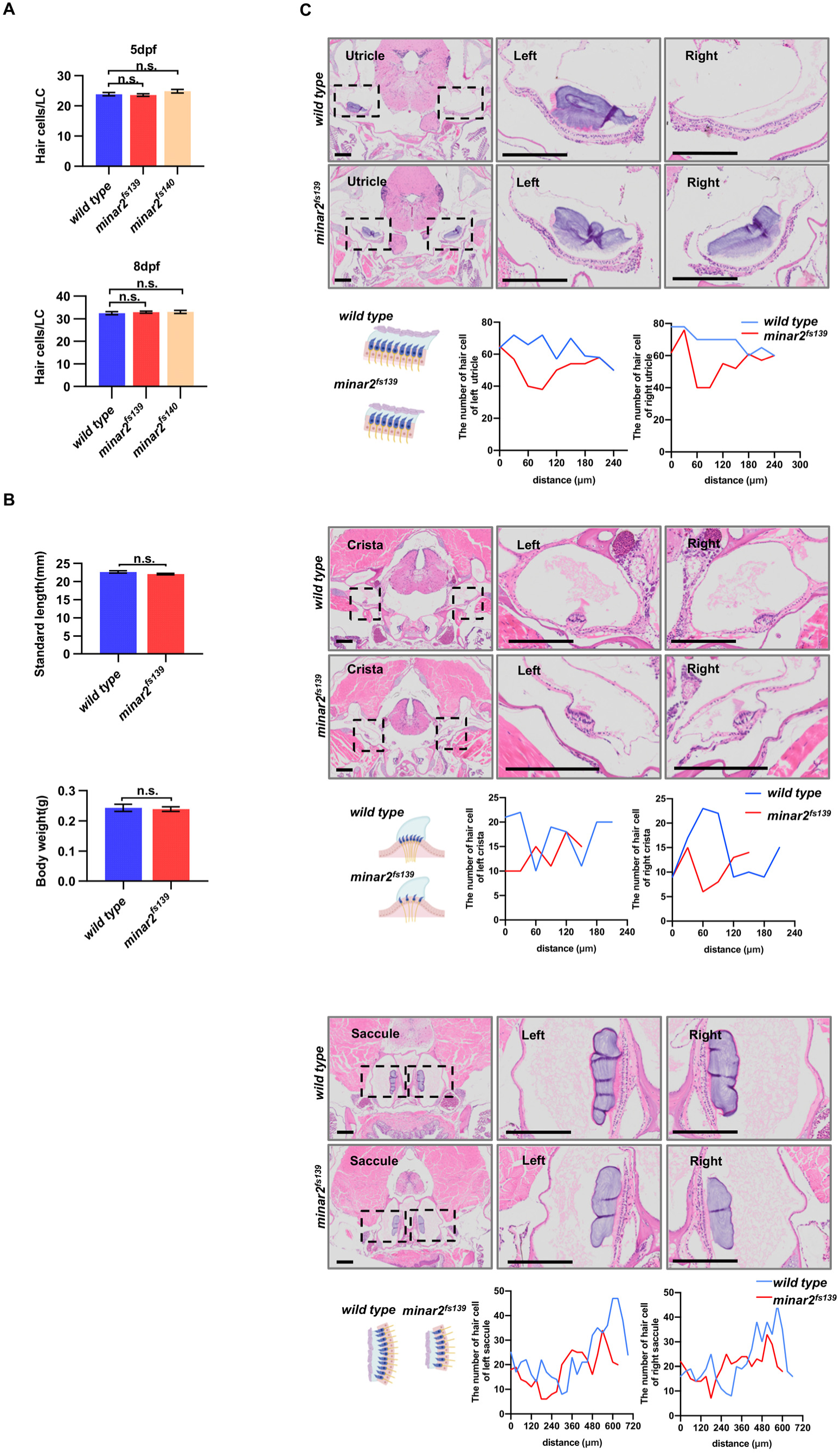
Numbers of inner ear hair cells in zebrafish larvae and adults. **(A)** Quantification of hair cell numbers in the inner ears of zebafish larvae. Hair bundles were labeled with fluorescence-conjugated phalloidin then the lateral crista regions were imaged and counted (for 5 dpf, n = 34, 29, and 22; for 8 dpf, n = 36, 23 and 23. n.s.: not significant). **(B)** Quantification of body length and body weight in wild type and mutant *minar2^fs139^* adult zebrafish (6 mpf, n = 9 and 11. n.s.: not significant). **(C)** Hematoxylin and eosin staining of head sections of wild type and mutant *minar2^fs139^* adult zebrafish (12 mpf). The head regions were cross-sectioned and sampled every 30 μm. The numbers of hair cells were counted and plotted along the anterior-posterior axis. The lengths of utricle, crista and saccule appeared smaller in the *minar2^fs139^* mutant. Scale bars represent 200 μm.

**Figure 2-figure supplement 1.**
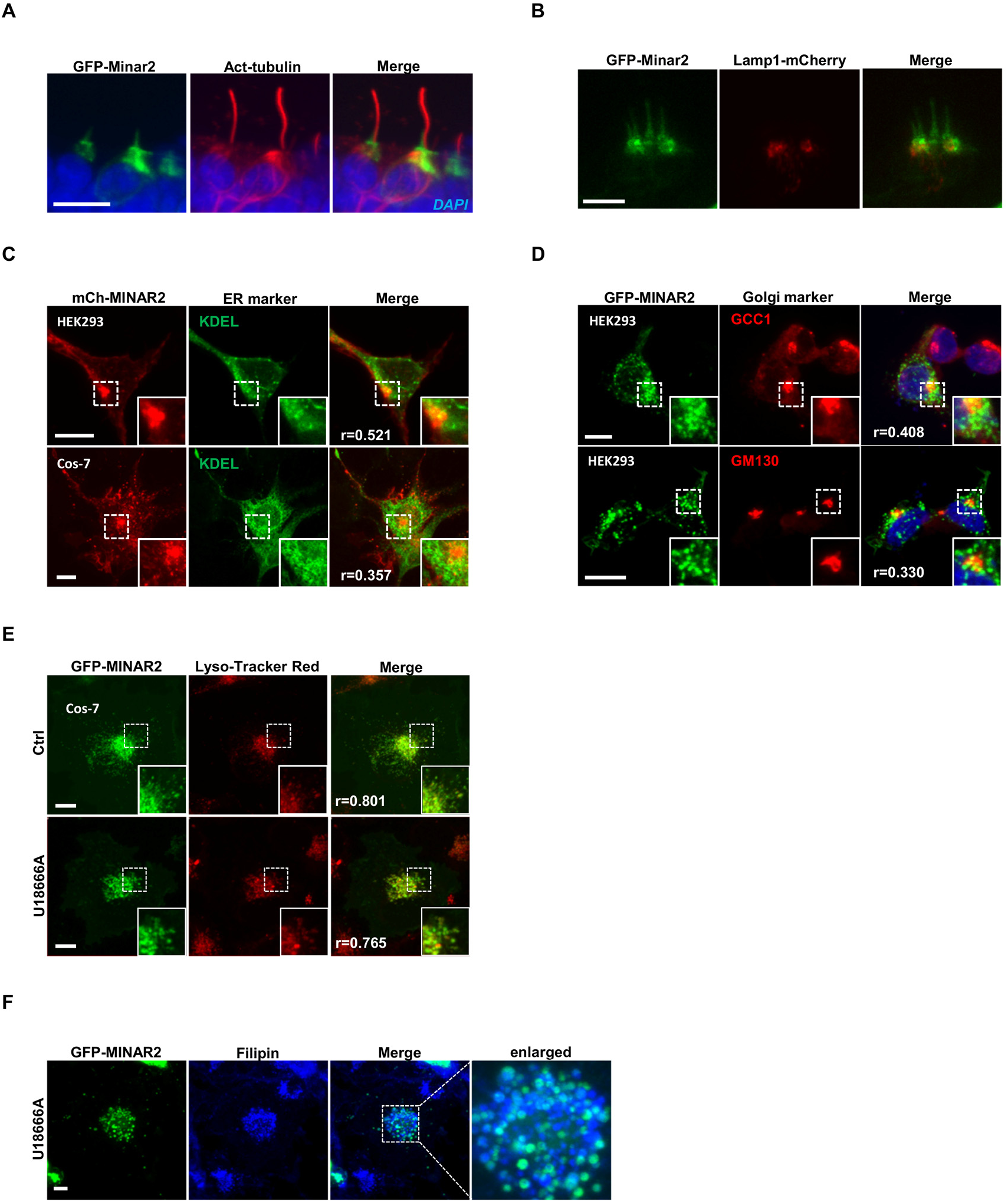
Subcellular localization of Minar2 protein. **(A)** Representative image of GFP-Minar2 localization in the inner ear hair cells. Kinocilia were stained with anti-acetylated tubulin antibodies (Act-tubulin). **(B)** Localization of GFP-Minar2 and Lamp1-mCherry in the inner ear hair cells. Representative image of lateral crista region. **(C-F)** Subcellular localization of GFP-Minar2 in cultured cells. Endoplasmic reticulum, Golgi complex, and lysosome are labeled by KDEL, GCC1/GM130, and Lyso-Tracker Red, respectively. Figure inserts show enlarged views of the boxed area. Pearson correlation coefficients (Pearson’s r values) are indicated. U18666A treatment traps cholesterol (stained by filipin) in the lysosome lumen. Scale bars represent 10 μm.

**Figure 4-figure supplement 1.**
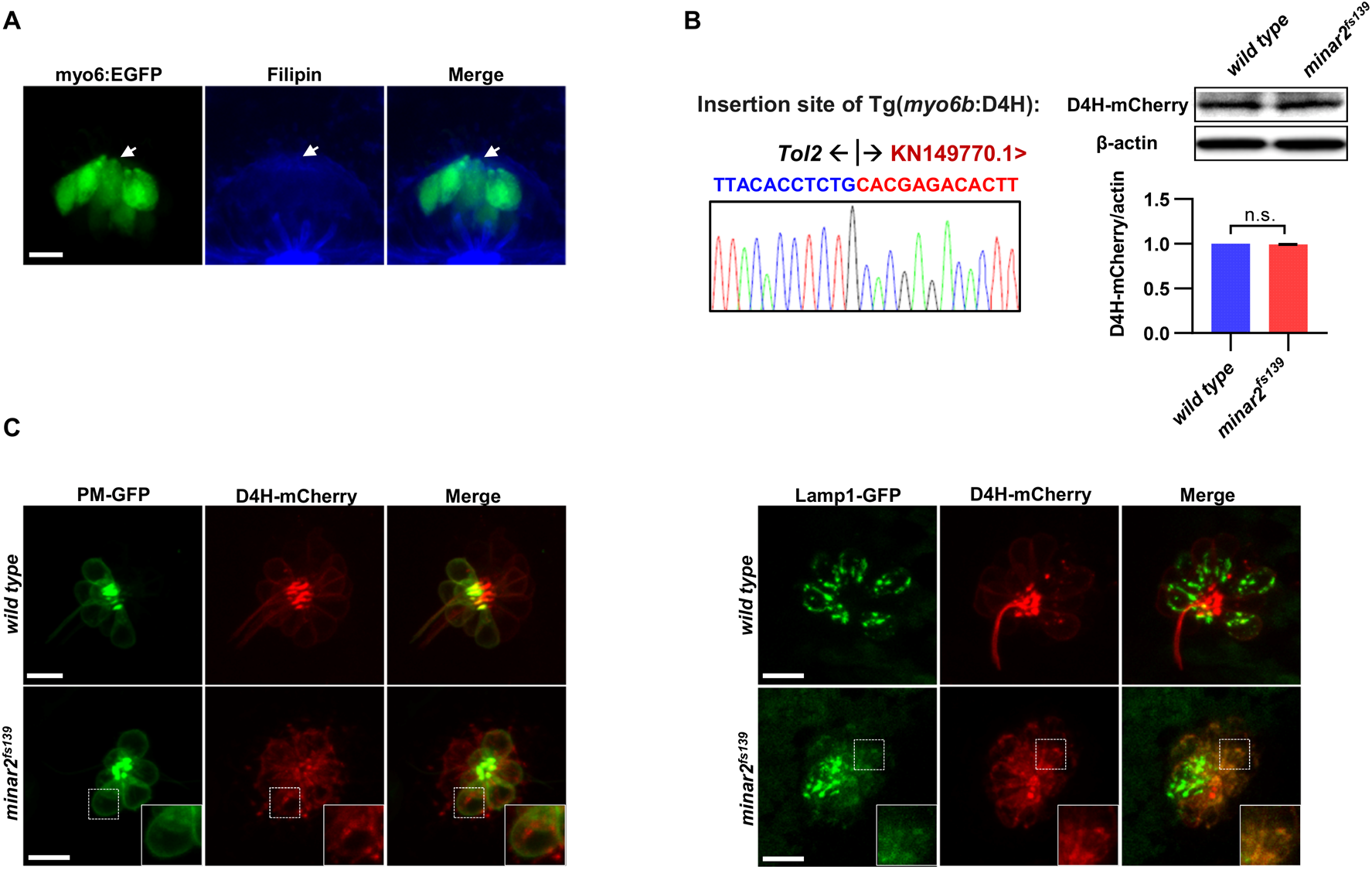
The abnormal vesicles in *minar2* mutant were co-labeled with lysosome. **(A)** Representative image of filipin staining of inner ear tissue. The lateral crista region of inner ear was imaged. The GFP signals in the *myo6:Gal4FF;UAS-EGFP* transgenic line (myo6:GFP) labels hair cells. White arrow points to the apical borders of hair cells. **(B)** Characterization of the Tg(*myo6b:*D4H-mCherry) report line. Genomic sequence at the transgenic insertion site is shown (left). Immunoblot and quantifications show that the expression levels of the D4H-mCherry transgene were not affected by the loss of *minar2* in zebrafish larvae. t = 1.25, df = 2, p = 0.337. **(C)** Representative images of D4H-mCherry labeled large vesicles in the lateral line hair cells and co-labeling by plasma membrane probe PM-GFP (left panels) and lysosomal marker Lamp1-GFP (right panels). D4H-mCherry labeled large vesicles were co-labeled by Lamp1-GFP. Figure inserts show enlarged views of the boxed area. Scale bars represent 10 μm.

**Figure 6-figure supplement 1.**
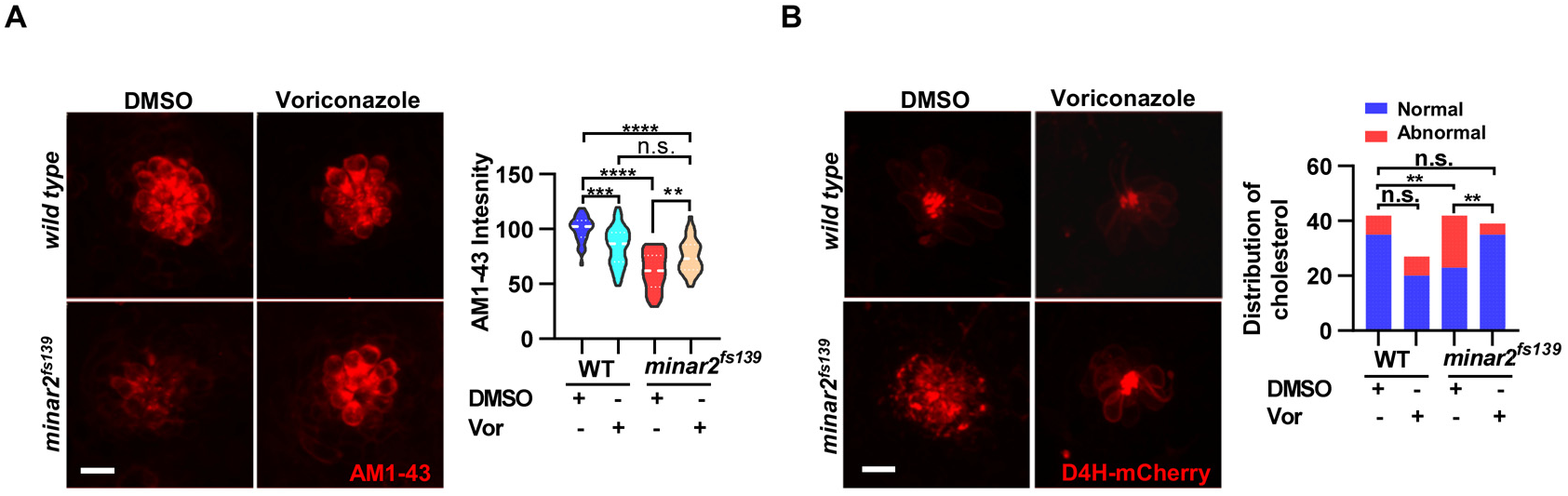
Increasing cholesterol levels by voriconazole rescue hair cell defects in *minar2* mutant. **(A)** Quantification of AM1-43 labeling in wild type and mutant *minar2^fs139^* larvae after voriconazole treatment. The lateral line L3 neuromast were imaged and quantified. n = 31, 43, 33, and 34. F(3, 131.4) = 34.50, p<0.0001. **(B)** Effects of voriconazole treatment on appearance of abnormally enlarged vesicles in hair cells of wild type and *minar2^fs139^* zebrafish. n = 42, 38, 42, and 34. χ^2^=20.92, df = 3, p<0.001. Multiple comparison significance values are indicated on the graph. Scale bars represent 10 μm.

**Figure 7-figure supplement 1.**
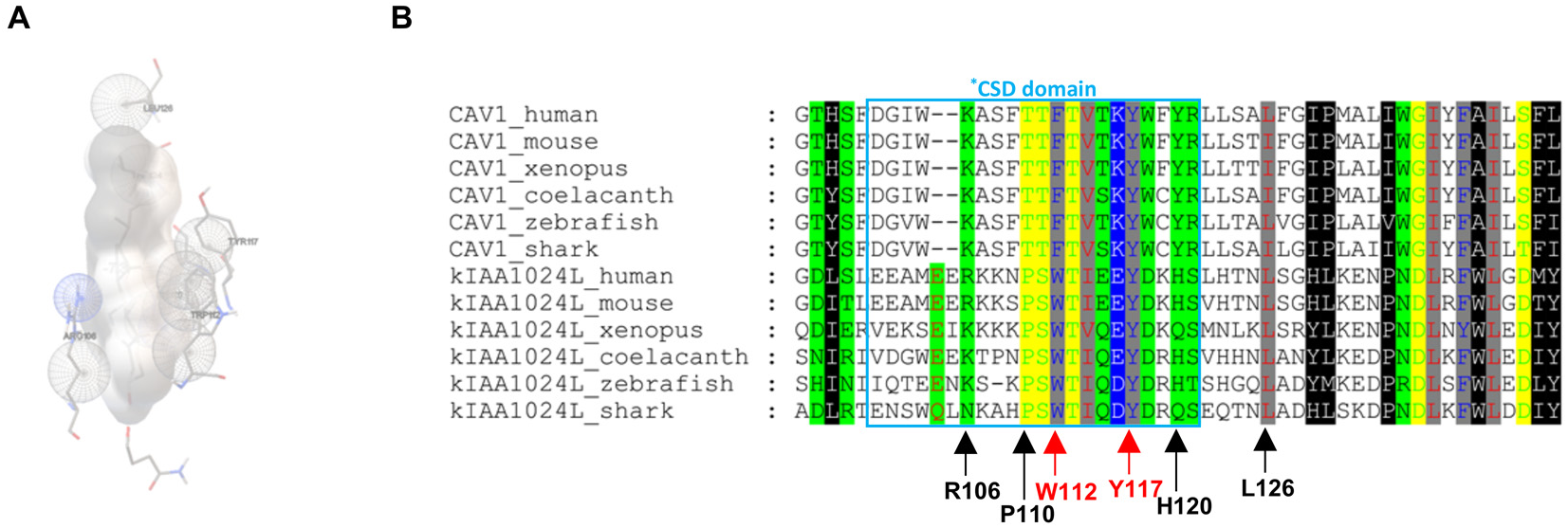
Computational docking between MINAR2 structure model and cholesterol. **(A)** Interaction between MINAR2 residues and cholesterol as modeled by AutoDock. MINAR2 residues in ball-and-stick view and cholesterol molecule in grey sphere view. **(B)** Cholesterol interacting residues of MINAR2 are conserved. Arrows point to residues showing interaction with cholesterol as modeled by AutoDock. The arrows pointing to the critical aromatic residues are labeled in red.

## References

Aasland, R., Abrams, C., Ampe, C., Ball, L. J., Bedford, M. T., Cesareni, G., Gimona, M., Hurley, J. H., Jarchau, T., Lehto, V. P., Lemmon, M. A., Linding, R., Mayer, B. J., Nagai, M., Sudol, M., Walter, U., & Winder, S. J. (2002). Normalization of nomenclature for peptide motifs as ligands of modular protein domains. FEBS Lett, 513(1), 141–144. https://doi.org/10.1016/s0014-5793(01)03295-1

Aqul, A., Liu, B., Ramirez, C. M., Pieper, A. A., Estill, S. J., Burns, D. K., Liu, B., Repa, J. J., Turley, S. D., & Dietschy, J. M. (2011). Unesterified cholesterol accumulation in late endosomes/lysosomes causes neurodegeneration and is prevented by driving cholesterol export from this compartment. J Neurosci, 31(25), 9404–9413. https://doi.org/10.1523/JNEUROSCI.1317-11.2011

Asakawa, K., Suster, M. L., Mizusawa, K., Nagayoshi, S., Kotani, T., Urasaki, A., Kishimoto, Y., Hibi, M., & Kawakami, K. (2008). Genetic dissection of neural circuits by Tol2 transposon-mediated Gal4 gene and enhancer trapping in zebrafish. Proc Natl Acad Sci U S A, 105(4), 1255–1260. https://doi.org/10.1073/pnas.0704963105

Atkinson, P. J., Huarcaya Najarro, E., Sayyid, Z. N., & Cheng, A. G. (2015). Sensory hair cell development and regeneration: similarities and differences. Development, 142(9), 1561–1571. https://doi.org/10.1242/dev.114926

Barr-Gillespie, P. G. (2015). Assembly of hair bundles, an amazing problem for cell biology. Mol Biol Cell, 26(15), 2727–2732. https://doi.org/10.1091/mbc.E14-04-0940

Barta, C. L., Liu, H., Chen, L., Giffen, K. P., Li, Y., Kramer, K. L., Beisel, K. W., & He, D. Z. (2018). RNA-seq transcriptomic analysis of adult zebrafish inner ear hair cells. Sci Data, 5, 180005. https://doi.org/10.1038/sdata.2018.5

Bartsch, T. F., Hengel, F. E., Oswald, A., Dionne, G., Chipendo, I. V., Mangat, S. S., El Shatanofy, M., Shapiro, L., Muller, U., & Hudspeth, A. J. (2019). Elasticity of individual protocadherin 15 molecules implicates tip links as the gating springs for hearing. Proc Natl Acad Sci U S A, 116(22), 11048–11056. https://doi.org/10.1073/pnas.1902163116

Basch, M. L., Brown, R. M., 2nd, Jen, H. I., & Groves, A. K. (2016). Where hearing starts: the development of the mammalian cochlea. J Anat, 228(2), 233–254. https://doi.org/10.1111/joa.12314

Belyantseva, I. A., Perrin, B. J., Sonnemann, K. J., Zhu, M., Stepanyan, R., McGee, J., Frolenkov, G. I., Walsh, E. J., Friderici, K. H., Friedman, T. B., & Ervasti, J. M. (2009). Gamma-actin is required for cytoskeletal maintenance but not development. Proc Natl Acad Sci U S A, 106(24), 9703–9708. https://doi.org/10.1073/pnas.0900221106

Blanco-Sanchez, B., Clement, A., Fierro, J., Jr., Washbourne, P., & Westerfield, M. (2014). Complexes of Usher proteins preassemble at the endoplasmic reticulum and are required for trafficking and ER homeostasis. Dis Model Mech, 7(5), 547–559. https://doi.org/10.1242/dmm.014068

Blanco-Sanchez, B., Clement, A., Phillips, J. B., & Westerfield, M. (2017). Zebrafish models of human eye and inner ear diseases. Methods Cell Biol, 138, 415–467. https://doi.org/10.1016/bs.mcb.2016.10.006

Bowl, M. R., Simon, M. M., Ingham, N. J., Greenaway, S., Santos, L., Cater, H., Taylor, S., Mason, J., Kurbatova, N., Pearson, S., Bower, L. R., Clary, D. A., Meziane, H., Reilly, P., Minowa, O., Kelsey, L., International Mouse Phenotyping, C., Tocchini-Valentini, G. P., Gao, X., . . . Brown, S. D. M. (2017). A large scale hearing loss screen reveals an extensive unexplored genetic landscape for auditory dysfunction. Nat Commun, 8(1), 886. https://doi.org/10.1038/s41467-017-00595-4

Burgess, H. A., & Granato, M. (2007). Sensorimotor gating in larval zebrafish. J Neurosci, 27(18), 4984–4994. https://doi.org/10.1523/JNEUROSCI.0615-07.2007

Buyan, A., Cox, C. D., Barnoud, J., Li, J., Chan, H. S. M., Martinac, B., Marrink, S. J., & Corry, B. (2020). Piezo1 Forms Specific, Functionally Important Interactions with Phosphoinositides and Cholesterol. Biophys J, 119(8), 1683–1697. https://doi.org/10.1016/j.bpj.2020.07.043

Chakraborty, S., Doktorova, M., Molugu, T. R., Heberle, F. A., Scott, H. L., Dzikovski, B., Nagao, M., Stingaciu, L. R., Standaert, R. F., Barrera, F. N., Katsaras, J., Khelashvili, G., Brown, M. F., & Ashkar, R. (2020). How cholesterol stiffens unsaturated lipid membranes. Proc Natl Acad Sci U S A, 117(36), 21896–21905. https://doi.org/10.1073/pnas.2004807117

Corwin, J. T., & Warchol, M. E. (1991). Auditory hair cells: structure, function, development, and regeneration. Annu. Rev. Neurosci., 14:301-33.

Crumling, M. A., Liu, L., Thomas, P. V., Benson, J., Kanicki, A., Kabara, L., Halsey, K., Dolan, D., & Duncan, R. K. (2012). Hearing loss and hair cell death in mice given the cholesterol-chelating agent hydroxypropyl-beta-cyclodextrin. PLoS One, 7(12), e53280. https://doi.org/10.1371/journal.pone.0053280

Cunningham, C. L., Qiu, X., Wu, Z., Zhao, B., Peng, G., Kim, Y. H., Lauer, A., & Muller, U. (2020). TMIE Defines Pore and Gating Properties of the Mechanotransduction Channel of Mammalian Cochlear Hair Cells. Neuron, 107(1), 126–143 e128. https://doi.org/10.1016/j.neuron.2020.03.033

Das, A., Brown, M. S., Anderson, D. D., Goldstein, J. L., & Radhakrishnan, A. (2014). Three pools of plasma membrane cholesterol and their relation to cholesterol homeostasis. Elife, 3. https://doi.org/10.7554/eLife.02882

Dimova, R. (2014). Recent developments in the field of bending rigidity measurements on membranes. Adv Colloid Interface Sci, 208, 225–234. https://doi.org/10.1016/j.cis.2014.03.003

Ding, D., Manohar, S., Jiang, H., & Salvi, R. (2020). Hydroxypropyl-beta-cyclodextrin causes massive damage to the developing auditory and vestibular system. Hear Res, 396, 108073. https://doi.org/10.1016/j.heares.2020.108073

Driver, E. C., & Kelley, M. W. (2020). Development of the cochlea. Development, 147(12). https://doi.org/10.1242/dev.162263

Effertz, T., Becker, L., Peng, A. W., & Ricci, A. J. (2017). Phosphoinositol-4,5-Bisphosphate Regulates Auditory Hair-Cell Mechanotransduction-Channel Pore Properties and Fast Adaptation. J Neurosci, 37(48), 11632–11646. https://doi.org/10.1523/JNEUROSCI.1351-17.2017

Elkon, R., Milon, B., Morrison, L., Shah, M., Vijayakumar, S., Racherla, M., Leitch, C. C., Silipino, L., Hadi, S., Weiss-Gayet, M., Barras, E., Schmid, C. D., Ait-Lounis, A., Barnes, A., Song, Y., Eisenman, D. J., Eliyahu, E., Frolenkov, G. I., Strome, S. E., . . . Hertzano, R. (2015). RFX transcription factors are essential for hearing in mice. Nat Commun, 6, 8549. https://doi.org/10.1038/ncomms9549

Epand, R. M., Sayer, B. G., & Epand, R. F. (2005). Caveolin scaffolding region and cholesterol-rich domains in membranes. J Mol Biol, 345(2), 339–350. https://doi.org/10.1016/j.jmb.2004.10.064

Erickson, T., & Nicolson, T. (2015). Identification of sensory hair-cell transcripts by thiouracil-tagging in zebrafish. BMC Genomics, 16, 842. https://doi.org/10.1186/s12864-015-2072-5

Everson, W. V., & Smart, E. J. (2005). Caveolae and the regulation of cellular cholesterol homeostasis. In M. Lisanti & P. Frank (Eds.), Caveolae and Lipid Rafts: Roles in Signal Transduction and the Pathogenesis of Human Diseases (Vol. 36, pp. 37–55). Elsevier. https://www.ncbi.nlm.nih.gov/pubmed/16603689

Fantini, J., Di Scala, C., Evans, L. S., Williamson, P. T., & Barrantes, F. J. (2016). A mirror code for protein-cholesterol interactions in the two leaflets of biological membranes. Sci Rep, 6, 21907. https://doi.org/10.1038/srep21907

Fettiplace, R. (2017). Hair Cell Transduction, Tuning, and Synaptic Transmission in the Mammalian Cochlea. Compr Physiol, 7(4), 1197–1227. https://doi.org/10.1002/cphy.c160049

Flock, A. (1964). Structure of the Macula Utriculi with Special Reference to Directional Interplay of Sensory Responses as Revealed by Morphological Polarization. J Cell Biol, 22, 413–431. https://doi.org/10.1083/jcb.22.2.413

Forge, A., Davies, S., & Zajic, G. (1988). Characteristics of the membrane of the stereocilia and cell apex in cochlear hair cells. J Neurocytol, 17(3), 325–334. https://doi.org/10.1007/bf01187855

Forge, A., & Richardson, G. (1993). Freeze fracture analysis of apical membranes in cochlear cultures: differences between basal and apical-coil outer hair cells and effects of neomycin. J Neurocytol, 22(10), 854–867. https://doi.org/10.1007/bf01186357

Forli, S., Huey, R., Pique, M. E., Sanner, M. F., Goodsell, D. S., & Olson, A. J. (2016). Computational protein-ligand docking and virtual drug screening with the AutoDock suite. Nat Protoc, 11(5), 905–919. https://doi.org/10.1038/nprot.2016.051

Frank, P. G., Cheung, M. W., Pavlides, S., Llaverias, G., Park, D. S., & Lisanti, M. P. (2006). Caveolin-1 and regulation of cellular cholesterol homeostasis. Am J Physiol Heart Circ Physiol, 291(2), H677–686. https://doi.org/10.1152/ajpheart.01092.2005

Glitscher, M., & Hildt, E. (2021). Endosomal Cholesterol in Viral Infections - A Common Denominator? Front Physiol, 12, 750544. https://doi.org/10.3389/fphys.2021.750544

Guo, Y., Zhang, C., Du, X., Nair, U., & Yoo, T. J. (2005). Morphological and functional alterations of the cochlea in apolipoprotein E gene deficient mice. Hear Res, 208(1-2), 54–67. https://doi.org/10.1016/j.heares.2005.05.010

Harris, J. R. (2010). Cholesterol binding and cholesterol transport proteins : structure and function in health and disease. Springer.

Higgs, D. M., Rollo, A. K., Souza, M. J., & Popper, A. N. (2003). Development of form and function in peripheral auditory structures of the zebrafish (Danio rerio). J Acoust Soc Am, 113(2), 1145–1154. https://doi.org/10.1121/1.1536185

Higgs, D. M., Souza, M. J., Wilkins, H. R., Presson, J. C., & Popper, A. N. (2002). Age- and size-related changes in the inner ear and hearing ability of the adult zebrafish (Danio rerio). J Assoc Res Otolaryngol, 3(2), 174–184. https://doi.org/10.1007/s101620020035

Hirono, M., Denis, C. S., Richardson, G. P., & Gillespie, P. G. (2004). Hair cells require phosphatidylinositol 4,5-bisphosphate for mechanical transduction and adaptation. Neuron, 44(2), 309–320. https://doi.org/10.1016/j.neuron.2004.09.020

Ho, R. X., Amraei, R., De La Cena, K. O. C., Sutherland, E. G., Mortazavi, F., Stein, T., Chitalia, V., & Rahimi, N. (2020). Loss of MINAR2 impairs motor function and causes Parkinson’s disease-like symptoms in mice. Brain Commun, 2(1), fcaa047. https://doi.org/10.1093/braincomms/fcaa047

Ho, R. X., Meyer, R. D., Chandler, K. B., Ersoy, E., Park, M., Bondzie, P. A., Rahimi, N., Xu, H., Costello, C. E., & Rahimi, N. (2018). MINAR1 is a Notch2-binding protein that inhibits angiogenesis and breast cancer growth. J Mol Cell Biol, 10(3), 195–204. https://doi.org/10.1093/jmcb/mjy002

Hudspeth, A. J. (1989). How the ear’s works work. Nature, 341(6241), 397–404. https://doi.org/10.1038/341397a0

Ikonen, E., Heino, S., & Lusa, S. (2004). Caveolins and membrane cholesterol. Biochem Soc Trans, 32(Pt 1), 121–123. https://doi.org/10.1042/bst0320121

Ingham, N. J., Pearson, S. A., Vancollie, V. E., Rook, V., Lewis, M. A., Chen, J., Buniello, A., Martelletti, E., Preite, L., Lam, C. C., Weiss, F. D., Powis, Z., Suwannarat, P., Lelliott, C. J., Dawson, S. J., White, J. K., & Steel, K. P. (2019). Mouse screen reveals multiple new genes underlying mouse and human hearing loss. PLoS Biol, 17(4), e3000194. https://doi.org/10.1371/journal.pbio.3000194

Kawakami, K., Shima, A., & Kawakami, N. (2000). Identification of a functional transposase of the Tol2 element, an Ac-like element from the Japanese medaka fish, and its transposition in the zebrafish germ lineage. Proc Natl Acad Sci U S A, 97(21), 11403–11408. https://doi.org/10.1073/pnas.97.21.11403

Kilsdonk, E. P., Yancey, P. G., Stoudt, G. W., Bangerter, F. W., Johnson, W. J., Phillips, M. C., & Rothblat, G. H. (1995). Cellular cholesterol efflux mediated by cyclodextrins. J Biol Chem, 270(29), 17250–17256. https://doi.org/10.1074/jbc.270.29.17250

Kindt, K. S., Finch, G., & Nicolson, T. (2012). Kinocilia mediate mechanosensitivity in developing zebrafish hair cells. Dev Cell, 23(2), 329–341. https://doi.org/10.1016/j.devcel.2012.05.022

King, K. A., Gordon-Salant, S., Pawlowski, K. S., Taylor, A. M., Griffith, A. J., Houser, A., Kurima, K., Wassif, C. A., Wright, C. G., Porter, F. D., Repa, J. J., & Brewer, C. C. (2014). Hearing loss is an early consequence of Npc1 gene deletion in the mouse model of Niemann-Pick disease, type C. J Assoc Res Otolaryngol, 15(4), 529–541. https://doi.org/10.1007/s10162-014-0459-7

King, K. A., Gordon-Salant, S., Yanjanin, N., Zalewski, C., Houser, A., Porter, F. D., & Brewer, C. C. (2014). Auditory phenotype of Niemann-Pick disease, type C1. Ear Hear, 35(1), 110–117. https://doi.org/10.1097/AUD.0b013e3182a362b8

Kneen, M., Farinas, J., Li, Y., & Verkman, A. S. (1998). Green fluorescent protein as a noninvasive intracellular pH indicator. Biophys J, 74(3), 1591–1599. https://doi.org/10.1016/S0006-3495(98)77870-1

Kozlov, A. S., Risler, T., & Hudspeth, A. J. (2007). Coherent motion of stereocilia assures the concerted gating of hair-cell transduction channels. Nat Neurosci, 10(1), 87–92. https://doi.org/10.1038/nn1818

Kratzel, A., Kelly, J. N., V’Kovski, P., Portmann, J., Bruggemann, Y., Todt, D., Ebert, N., Shrestha, N., Plattet, P., Staab-Weijnitz, C. A., von Brunn, A., Steinmann, E., Dijkman, R., Zimmer, G., Pfaender, S., & Thiel, V. (2021). A genome-wide CRISPR screen identifies interactors of the autophagy pathway as conserved coronavirus targets. PLoS Biol, 19(12), e3001490. https://doi.org/10.1371/journal.pbio.3001490

Krey, J. F., Dumont, R. A., Wilmarth, P. A., David, L. L., Johnson, K. R., & Barr-Gillespie, P. G. (2018). ELMOD1 Stimulates ARF6-GTP Hydrolysis to Stabilize Apical Structures in Developing Vestibular Hair Cells. J Neurosci, 38(4), 843–857. https://doi.org/10.1523/JNEUROSCI.2658-17.2017

Lange, Y., & Steck, T. L. (2016). Active membrane cholesterol as a physiological effector. Chem Phys Lipids, 199, 74–93. https://doi.org/10.1016/j.chemphyslip.2016.02.003

Lange, Y., Ye, J., & Steck, T. L. (2004). How cholesterol homeostasis is regulated by plasma membrane cholesterol in excess of phospholipids. Proc Natl Acad Sci U S A, 101(32), 11664–11667. https://doi.org/10.1073/pnas.0404766101

Le Lan, C., Gallay, J., Vincent, M., Neumann, J. M., de Foresta, B., & Jamin, N. (2010). Structural and dynamic properties of juxta-membrane segments of caveolin-1 and caveolin-2 at the membrane interface. Eur Biophys J, 39(2), 307–325. https://doi.org/10.1007/s00249-009-0548-4

Li, H., & Papadopoulos, V. (1998). Peripheral-type benzodiazepine receptor function in cholesterol transport. Identification of a putative cholesterol recognition/interaction amino acid sequence and consensus pattern. Endocrinology, 139(12), 4991–4997. https://doi.org/10.1210/endo.139.12.6390

Liang, J., & Burgess, S. M. (2009). Gross and fine dissection of inner ear sensory epithelia in adult zebrafish (Danio rerio). J Vis Exp(27). https://doi.org/10.3791/1211

Lim, C. Y., Davis, O. B., Shin, H. R., Zhang, J., Berdan, C. A., Jiang, X., Counihan, J. L., Ory, D. S., Nomura, D. K., & Zoncu, R. (2019). ER-lysosome contacts enable cholesterol sensing by mTORC1 and drive aberrant growth signalling in Niemann-Pick type C. Nat Cell Biol, 21(10), 1206–1218. https://doi.org/10.1038/s41556-019-0391-5

Linkert, M., Rueden, C. T., Allan, C., Burel, J. M., Moore, W., Patterson, A., Loranger, B., Moore, J., Neves, C., Macdonald, D., Tarkowska, A., Sticco, C., Hill, E., Rossner, M., Eliceiri, K. W., & Swedlow, J. R. (2010). Metadata matters: access to image data in the real world. J Cell Biol, 189(5), 777–782. https://doi.org/10.1083/jcb.201004104

Liu, B., Turley, S. D., Burns, D. K., Miller, A. M., Repa, J. J., & Dietschy, J. M. (2009). Reversal of defective lysosomal transport in NPC disease ameliorates liver dysfunction and neurodegeneration in the npc1-/- mouse. Proc Natl Acad Sci U S A, 106(7), 2377–2382. https://doi.org/10.1073/pnas.0810895106

Liu, F., Baggerman, G., Schoofs, L., & Wets, G. (2006). Uncovering conserved patterns in bioactive peptides in Metazoa. Peptides, 27(12), 3137–3153. https://doi.org/10.1016/j.peptides.2006.08.021

Liu, H., Chen, L., Giffen, K. P., Stringham, S. T., Li, Y., Judge, P. D., Beisel, K. W., & He, D. Z. Z. (2018). Cell-Specific Transcriptome Analysis Shows That Adult Pillar and Deiters’ Cells Express Genes Encoding Machinery for Specializations of Cochlear Hair Cells. Front Mol Neurosci, 11, 356. https://doi.org/10.3389/fnmol.2018.00356

Liu, H., Yang, L., Zhang, Q., Mao, L., Jiang, H., & Yang, H. (2016). Probing the structure and dynamics of caveolin-1 in a caveolae-mimicking asymmetric lipid bilayer model. Eur Biophys J, 45(6), 511–521. https://doi.org/10.1007/s00249-016-1118-1

Liu, Y. G., & Chen, Y. (2007). High-efficiency thermal asymmetric interlaced PCR for amplification of unknown flanking sequences. Biotechniques, 43(5), 649–650, 652, 654 passim. https://doi.org/10.2144/000112601

Liu, Y. G., & Whittier, R. F. (1995). Thermal asymmetric interlaced PCR: automatable amplification and sequencing of insert end fragments from P1 and YAC clones for chromosome walking. Genomics, 25(3), 674–681. https://doi.org/10.1016/0888-7543(95)80010-j

Livingstone, C. D., & Barton, G. J. (1993). Protein sequence alignments: a strategy for the hierarchical analysis of residue conservation. Comput Appl Biosci, 9(6), 745–756. https://doi.org/10.1093/bioinformatics/9.6.745

Lu, F., Liang, Q., Abi-Mosleh, L., Das, A., De Brabander, J. K., Goldstein, J. L., & Brown, M. S. (2015). Identification of NPC1 as the target of U18666A, an inhibitor of lysosomal cholesterol export and Ebola infection. Elife, 4. https://doi.org/10.7554/eLife.12177

Maeda, R., Pacentine, I. V., Erickson, T., & Nicolson, T. (2017). Functional Analysis of the Transmembrane and Cytoplasmic Domains of Pcdh15a in Zebrafish Hair Cells. J Neurosci, 37(12), 3231–3245. https://doi.org/10.1523/JNEUROSCI.2216-16.2017

Maekawa, M., & Fairn, G. D. (2015). Complementary probes reveal that phosphatidylserine is required for the proper transbilayer distribution of cholesterol. J Cell Sci, 128(7), 1422–1433. https://doi.org/10.1242/jcs.164715

Maxfield, F. R., & van Meer, G. (2010). Cholesterol, the central lipid of mammalian cells. Curr Opin Cell Biol, 22(4), 422–429. https://doi.org/10.1016/j.ceb.2010.05.004

Maxfield, F. R., & Wustner, D. (2012). Analysis of cholesterol trafficking with fluorescent probes. Methods Cell Biol, 108, 367–393. https://doi.org/10.1016/B978-0-12-386487-1.00017-1

McGrath, J., Roy, P., & Perrin, B. J. (2017). Stereocilia morphogenesis and maintenance through regulation of actin stability. Semin Cell Dev Biol, 65, 88–95. https://doi.org/10.1016/j.semcdb.2016.08.017

Meyers, J. R., MacDonald, R. B., Duggan, A., Lenzi, D., Standaert, D. G., Corwin, J. T., & Corey, D. P. (2003). Lighting up the Senses: FM1-43 Loading of Sensory Cells through Nonselective Ion Channels. The Journal of Neuroscience, 23(10), 4054–4065. https://doi.org/10.1523/jneurosci.23-10-04054.2003

Monroe, J. D., Manning, D. P., Uribe, P. M., Bhandiwad, A., Sisneros, J. A., Smith, M. E., & Coffin, A. B. (2016). Hearing sensitivity differs between zebrafish lines used in auditory research. Hear Res, 341, 220–231. https://doi.org/10.1016/j.heares.2016.09.004

Morizono, T., and Paparella, M. M. (1978). Hypercholesterolemia and Auditory Dysfunction: Experimental Studies. Ann. Otol. Rhinol. Laryngol., 87,804–814.

Morris, G. M., Huey, R., Lindstrom, W., Sanner, M. F., Belew, R. K., Goodsell, D. S., & Olson, A. J. (2009). AutoDock4 and AutoDockTools4: Automated docking with selective receptor flexibility. J Comput Chem, 30(16), 2785–2791. https://doi.org/10.1002/jcc.21256

Murata, M., Peränen, J., Schreiner, R., Wieland, F., Kurzchalia, T. V., & Simons, K. (1995). VIP21/caveolin is a cholesterol-binding protein. Proc Natl Acad Sci U S A, 92(22), 10339–10343. https://doi.org/10.1073/pnas.92.22.10339

Naito, T., Ercan, B., Krshnan, L., Triebl, A., Koh, D. H. Z., Wei, F. Y., Tomizawa, K., Torta, F. T., Wenk, M. R., & Saheki, Y. (2019). Movement of accessible plasma membrane cholesterol by the GRAMD1 lipid transfer protein complex. Elife, 8. https://doi.org/10.7554/eLife.51401

Nguyen, T. V., & Brownell, W. E. (1998). Contribution of membrane cholesterol to outer hair cell lateral wall stiffness. Otolaryngol Head Neck Surg, 119(1), 14–20. https://doi.org/10.1016/s0194-5998(98)70167-6

Noben-Trauth, K., Zheng, Q. Y., & Johnson, K. R. (2003). Association of cadherin 23 with polygenic inheritance and genetic modification of sensorineural hearing loss. Nat Genet, 35(1), 21–23. https://doi.org/10.1038/ng1226

O. Maoileidigh, D., & Ricci, A. J. (2019). A Bundle of Mechanisms: Inner-Ear Hair-Cell Mechanotransduction. Trends Neurosci, 42(3), 221–236. https://doi.org/10.1016/j.tins.2018.12.006

Paschkowsky, S., Recinto, S. J., Young, J. C., Bondar, A. N., & Munter, L. M. (2018). Membrane cholesterol as regulator of human rhomboid protease RHBDL4. J Biol Chem, 293(40), 15556–15568. https://doi.org/10.1074/jbc.RA118.002640

Paul, D., Hoppe, S., Saher, G., Krijnse-Locker, J., & Bartenschlager, R. (2013). Morphological and biochemical characterization of the membranous hepatitis C virus replication compartment. J Virol, 87(19), 10612–10627. https://doi.org/10.1128/jvi.01370-13

Perrin, B. J., Strandjord, D. M., Narayanan, P., Henderson, D. M., Johnson, K. R., & Ervasti, J. M. (2013). beta-Actin and fascin-2 cooperate to maintain stereocilia length. J Neurosci, 33(19), 8114–8121. https://doi.org/10.1523/JNEUROSCI.0238-13.2013

Petrov, A. M., & Pikuleva, I. A. (2019). Cholesterol 24-Hydroxylation by CYP46A1: Benefits of Modulation for Brain Diseases. Neurotherapeutics, 16(3), 635–648. https://doi.org/10.1007/s13311-019-00731-6

Powers, R. J., Kulason, S., Atilgan, E., Brownell, W. E., Sun, S. X., Barr-Gillespie, P. G., & Spector, A. A. (2014). The local forces acting on the mechanotransduction channel in hair cell stereocilia. Biophys J, 106(11), 2519–2528. https://doi.org/10.1016/j.bpj.2014.03.034

Powers, R. J., Roy, S., Atilgan, E., Brownell, W. E., Sun, S. X., Gillespie, P. G., & Spector, A. A. (2012). Stereocilia membrane deformation: implications for the gating spring and mechanotransduction channel. Biophys J, 102(2), 201–210. https://doi.org/10.1016/j.bpj.2011.12.022

Purcell, E. K., Liu, L., Thomas, P. V., & Duncan, R. K. (2011). Cholesterol influences voltage-gated calcium channels and BK-type potassium channels in auditory hair cells. PLoS One, 6(10), e26289. https://doi.org/10.1371/journal.pone.0026289

Pyenta, P. S., Holowka, D., & Baird, B. (2001). Cross-correlation analysis of inner-leaflet-anchored green fluorescent protein co-redistributed with IgE receptors and outer leaflet lipid raft components. Biophys J, 80(5), 2120–2132. https://doi.org/10.1016/s0006-3495(01)76185-1

Rajagopalan, L., Greeson, J. N., Xia, A., Liu, H., Sturm, A., Raphael, R. M., Davidson, A. L., Oghalai, J. S., Pereira, F. A., & Brownell, W. E. (2007). Tuning of the outer hair cell motor by membrane cholesterol. J Biol Chem, 282(50), 36659–36670. https://doi.org/10.1074/jbc.M705078200

Razani, B., Woodman, S. E., & Lisanti, M. P. (2002). Caveolae: from cell biology to animal physiology. Pharmacol Rev, 54(3), 431–467. https://doi.org/10.1124/pr.54.3.431

Revelo, N. H., Kamin, D., Truckenbrodt, S., Wong, A. B., Reuter-Jessen, K., Reisinger, E., Moser, T., & Rizzoli, S. O. (2014). A new probe for super-resolution imaging of membranes elucidates trafficking pathways. J Cell Biol, 205(4), 591–606. https://doi.org/10.1083/jcb.201402066

Ridone, P., Pandzic, E., Vassalli, M., Cox, C. D., Macmillan, A., Gottlieb, P. A., & Martinac, B. (2020). Disruption of membrane cholesterol organization impairs the activity of PIEZO1 channel clusters. J Gen Physiol, 152(8). https://doi.org/10.1085/jgp.201912515

Sasavage, N. L., Nilson, J. H., Horowitz, S., & Rottman, F. M. (1982). Nucleotide sequence of bovine prolactin messenger RNA. Evidence for sequence polymorphism. J Biol Chem, 257(2), 678–681. https://www.ncbi.nlm.nih.gov/pubmed/6274859

Schlegel, A., Schwab, R. B., Scherer, P. E., & Lisanti, M. P. (1999). A role for the caveolin scaffolding domain in mediating the membrane attachment of caveolin-1. The caveolin scaffolding domain is both necessary and sufficient for membrane binding in vitro. J Biol Chem, 274(32), 22660–22667. https://doi.org/10.1074/jbc.274.32.22660

Schneider, C. A., Rasband, W. S., & Eliceiri, K. W. (2012). NIH Image to ImageJ: 25 years of image analysis. Nat Methods, 9(7), 671–675. https://doi.org/10.1038/nmeth.2089

Schoop, V., Martello, A., Eden, E. R., & Hoglinger, D. (2021). Cellular cholesterol and how to find it. Biochim Biophys Acta Mol Cell Biol Lipids, 1866(9), 158989. https://doi.org/10.1016/j.bbalip.2021.158989

Severs, N. J. (1997). Cholesterol Cytochemistry in Cell Biology and Disease. In R. Bittman (Ed.), Cholesterol: Its Functions and Metabolism in Biology and Medicine (pp. 477–506). Springer.

Shearer, A. E., Hildebrand, M. S., & Smith, R. J. H. (1993). Hereditary Hearing Loss and Deafness Overview. In M. P. Adam, H. H. Ardinger, R. A. Pagon, S. E. Wallace, L. J. H. Bean, K. W. Gripp, G. M. Mirzaa, & A. Amemiya (Eds.), GeneReviews(®). University of Washington, Seattle Copyright © 1993-2022, University of Washington, Seattle. GeneReviews is a registered trademark of the University of Washington, Seattle. All rights reserved.

Sikora, M. A., Morizono, T., Ward, W. D., Paparella, M. M., & Leslie, K. (1986). Diet-induced hyperlipidemia and auditory dysfunction. Acta Otolaryngol, 102(5-6), 372–381. https://doi.org/10.3109/00016488609119420

Spicer, S. S., Thomopoulos, G. N., & Schulte, B. A. (1999). Novel membranous structures in apical and basal compartments of inner hair cells. J Comp Neurol, 409(3), 424–437.

Steinhart, M. R., Serdy, S. A., van der Valk, W. H., Zhang, J., Kim, J., Lee, J., & Koehler, K. R. (2022). Defining inner ear cell type specification at single-cell resolution in a model of human cranial development. Available at SSRN: https://ssrn.com/abstract=3974124 or http://dx.doi.org/10.2139/ssrn.3974124.

Stoeck, I. K., Lee, J. Y., Tabata, K., Romero-Brey, I., Paul, D., Schult, P., Lohmann, V., Kaderali, L., & Bartenschlager, R. (2018). Hepatitis C Virus Replication Depends on Endosomal Cholesterol Homeostasis. J Virol, 92(1). https://doi.org/10.1128/JVI.01196-17

Subczynski, W. K., Pasenkiewicz-Gierula, M., Widomska, J., Mainali, L., & Raguz, M. (2017). High Cholesterol/Low Cholesterol: Effects in Biological Membranes: A Review. Cell Biochem Biophys, 75(3-4), 369–385. https://doi.org/10.1007/s12013-017-0792-7

Takahashi, S., Homma, K., Zhou, Y., Nishimura, S., Duan, C., Chen, J., Ahmad, A., Cheatham, M. A., & Zheng, J. (2016). Susceptibility of outer hair cells to cholesterol chelator 2-hydroxypropyl-beta-cyclodextrine is prestin-dependent. Sci Rep, 6, 21973. https://doi.org/10.1038/srep21973

Thoenes, M., Zimmermann, U., Ebermann, I., Ptok, M., Lewis, M. A., Thiele, H., Morlot, S., Hess, M. M., Gal, A., Eisenberger, T., Bergmann, C., Nurnberg, G., Nurnberg, P., Steel, K. P., Knipper, M., & Bolz, H. J. (2015). OSBPL2 encodes a protein of inner and outer hair cell stereocilia and is mutated in autosomal dominant hearing loss (DFNA67). Orphanet J Rare Dis, 10, 15. https://doi.org/10.1186/s13023-015-0238-5

Tilney, L. G. (1980). The organization of actin filaments in the stereocilia of cochlear hair cells. J. Cell Biol., 86, 244–259.

Tilney, L. G., Tilney, M. S., & DeRosier, D. J. (1992). Actin filaments, stereocilia, and hair cells: how cells count and measure. Annu Rev Cell Biol, 8, 257–274. https://doi.org/10.1146/annurev.cb.08.110192.001353

Tunyasuvunakool, K., Adler, J., Wu, Z., Green, T., Zielinski, M., Žídek, A., Bridgland, A., Cowie, A., Meyer, C., Laydon, A., Velankar, S., Kleywegt, G. J., Bateman, A., Evans, R., Pritzel, A., Figurnov, M., Ronneberger, O., Bates, R., Kohl, S. A. A., . . . Hassabis, D. (2021). Highly accurate protein structure prediction for the human proteome. Nature, 596(7873), 590–596. https://doi.org/10.1038/s41586-021-03828-1

Twu, W. I., Lee, J. Y., Kim, H., Prasad, V., Cerikan, B., Haselmann, U., Tabata, K., & Bartenschlager, R. (2021). Contribution of autophagy machinery factors to HCV and SARS-CoV-2 replication organelle formation. Cell Rep, 37(8), 110049. https://doi.org/10.1016/j.celrep.2021.110049

Urasaki, A., Morvan, G., & Kawakami, K. (2006). Functional dissection of the Tol2 transposable element identified the minimal cis-sequence and a highly repetitive sequence in the subterminal region essential for transposition. Genetics, 174(2), 639–649. https://doi.org/10.1534/genetics.106.060244

Van Camp, G., & Smith, R. J. H. (2021). Hereditary Hearing Loss Homepage. https://hereditaryhearingloss.org https://hereditaryhearingloss.org

Wang, J., Song, Q., Yu, D., Yang, G., Xia, L., Su, K., Shi, H., Wang, J., & Yin, S. (2015). Ontogenetic development of the auditory sensory organ in zebrafish (Danio rerio): changes in hearing sensitivity and related morphology. Sci Rep, 5, 15943. https://doi.org/10.1038/srep15943

Wang, T., Zhou, M., Zhang, Q., Zhang, C., & Peng, G. (2021). ubtor Mutation Causes Motor Hyperactivity by Activating mTOR Signaling in Zebrafish. Neurosci Bull, 37(12), 1658–1670. https://doi.org/10.1007/s12264-021-00755-z

Westerfield, M. (2007). The Zebrafish Book: A guide for the laboratory use of zebrafish Danio rerio. University of Oregon Press.

Whitfield, T. T., Granato, M., van Eeden, F. J., Schach, U., Brand, M., Furutani-Seiki, M., Haffter, P., Hammerschmidt, M., Heisenberg, C. P., Jiang, Y. J., Kane, D. A., Kelsh, R. N., Mullins, M. C., Odenthal, J., & Nüsslein-Volhard, C. (1996). Mutations affecting development of the zebrafish inner ear and lateral line. Development, 123, 241–254. https://doi.org/10.1242/dev.123.1.241

Wiwatpanit, T., Remis, N. N., Ahmad, A., Zhou, Y., Clancy, J. C., Cheatham, M. A., & Garcia-Anoveros, J. (2018). Codeficiency of Lysosomal Mucolipins 3 and 1 in Cochlear Hair Cells Diminishes Outer Hair Cell Longevity and Accelerates Age-Related Hearing Loss. J Neurosci, 38(13), 3177–3189. https://doi.org/10.1523/JNEUROSCI.3368-17.2018

Wolman, M. A., Jain, R. A., Liss, L., & Granato, M. (2011). Chemical modulation of memory formation in larval zebrafish. Proc Natl Acad Sci U S A, 108(37), 15468–15473. https://doi.org/10.1073/pnas.1107156108

World-Health-Organization. (2021). World report on hearing

Wu, M., Holowka, D., Craighead, H. G., & Baird, B. (2004). Visualization of plasma membrane compartmentalization with patterned lipid bilayers. Proc Natl Acad Sci U S A, 101(38), 13798–13803. https://doi.org/10.1073/pnas.0403835101

Xing, G., Yao, J., Wu, B., Liu, T., Wei, Q., Liu, C., Lu, Y., Chen, Z., Zheng, H., Yang, X., & Cao, X. (2015). Identification of OSBPL2 as a novel candidate gene for progressive nonsyndromic hearing loss by whole-exome sequencing. Genet Med, 17(3), 210–218. https://doi.org/10.1038/gim.2014.90

Yao, J., Zeng, H., Zhang, M., Wei, Q., Wang, Y., Yang, H., Lu, Y., Li, R., Xiong, Q., Zhang, L., Chen, Z., Xing, G., Cao, X., & Dai, Y. (2019). OSBPL2-disrupted pigs recapitulate dual features of human hearing loss and hypercholesterolaemia. J Genet Genomics, 46(8), 379–387. https://doi.org/10.1016/j.jgg.2019.06.006

Zhang, H., Zhang, Q., Gao, G., Wang, X., Wang, T., Kong, Z., Wang, G., Zhang, C., Wang, Y., & Peng, G. (2018). UBTOR/KIAA1024 regulates neurite outgrowth and neoplasia through mTOR signaling. PLoS Genet, 14(8), e1007583. https://doi.org/10.1371/journal.pgen.1007583

Zidovetzki, R., & Levitan, I. (2007). Use of cyclodextrins to manipulate plasma membrane cholesterol content: evidence, misconceptions and control strategies. Biochim Biophys Acta, 1768(6), 1311–1324. https://doi.org/10.1016/j.bbamem.2007.03.026

